# Autotaxin facilitates selective LPA receptor signaling

**DOI:** 10.1101/2022.04.09.487723

**Authors:** Fernando Salgado-Polo, Razvan Borza, Florence Marsais, Catherine Jagerschmidt, Ludovic Waeckel, Wouter H. Moolenaar, Paul Ford, Bertrand Heckmann, Anastassis Perrakis

## Abstract

Autotaxin (ATX; ENPP2) produces the lipid mediator lysophosphatidic acid (LPA) that signals through disparate EDG (LPA_1-3_) and P2Y (LPA_4-6_) G protein-coupled receptors. ATX/LPA promote several (patho)physiological processes, including in pulmonary fibrosis, thus serving as attractive drug targets. However, it remains unclear if clinical outcome depends on how different ATX inhibitors modulate the ATX/LPA signaling axis. Here, we show that inhibitors binding to the ATX “tunnel” specifically abrogate key aspects of ATX/LPA signaling. We find that the tunnel is essential for signaling efficacy and dictates cellular responses independent of ATX catalytic activity, with a preference for activation of P2Y LPA receptors. These responses are abrogated by tunnel-binding inhibitors, such as ziritaxestat, but not by inhibitors that exclusively target the active site, as shown in primary lung fibroblasts and a murine model of radiation-induced pulmonary fibrosis. Our results uncover a receptor-selective signaling mechanism for ATX, implying clinical benefit for tunnel-targeting ATX inhibitors.

**Highlights:** ATX is a dual-function protein acting as an LPA-producing enzyme and LPA chaperone.

Structural integrity of the ATX tunnel is essential to mediate signaling functions.

ATX-bound LPA signals preferentially via P2Y family LPA receptors.

Occupancy of the ATX tunnel is crucial for ziritaxestat to exert inhibition *in vivo*.

## INTRODUCTION

Lysophosphatidic acid (LPA; mono-acyl-sn-glycero-3-phosphate) is a lipid mediator that signals through specific G protein-coupled receptors (GPCRs) to regulate multiple biological processes (Yanagida et al., 2013; Yung et al., 2014). Six LPA receptors (LPARs), belonging to two unrelated families, have been identified. LPA_1-3_ belong to the so-called EDG family, together with the receptors for sphingosine-1-phosphate (S1PR1-5), whereas LPA_4-6_ are P2Y (purinergic-type) GPCRs (Blaho and Chun, 2018). Strikingly, structural studies have revealed different modes of ligand (LPA) entry for the prototypic receptor, LPA_1_ (EDG2) versus that for LPA_6_ (P2Y5). LPA_1_ accepts its LPA ligand from the extracellular space (water phase), whereas LPA_6_ is thought to receive LPA by lateral diffusion, following insertion into the outer lipid bilayer (Chrencik et al., 2015; Taniguchi et al., 2017).

All LPARs activate diverse effector pathways by coupling to distinct heterotrimeric G proteins. The LPAR expression pattern and G protein coupling repertoire in a given cell type largely determine the signaling output, which includes proliferation and survival via AKT and ERK pathways, and migration and invasion via Rho-family small GTPases (Kihara et al., 2014). LPA signaling is restrained by the action of cell-associated lipid phosphate phosphatases (LPP1-3) (Morris and Smyth, 2014) that dephosphorylate LPA to non-signaling mono-acylglycerol.

LPA is produced by autotaxin (ATX or ENPP2), a secreted lyso-phospholipase D (lysoPLD) that hydrolyzes extracellular lysophosphatidylcholine (LPC) and other lysophospholipids into their bioactive products. ATX is secreted by diverse cell types and is present in body fluids; it is the only member of the ectonucleotide pyrophosphatase/phosphodiesterase family (ENPP) with lysoPLD activity (Borza et al., 2022). The ATX–LPAR signaling axis regulates numerous biological activities, including embryonic development and postnatal organ function (van Meeteren et al., 2006; Yasuda et al., 2019), and has been implicated in life-threatening diseases, particularly pulmonary fibrosis and cancer(Kaffe et al., 2019; Ninou et al., 2018). For this reason, ATX has attracted considerable interest as a drug target.

ATX is a compact, multi-domain glycoprotein consisting of a central bimetallic catalytic phosphodiesterase (PDE) domain flanked by two N□terminal somatomedin B-like domains and a C-terminal inactive nuclease-like domain (Hausmann et al., 2011; Nishimasu et al., 2011). The PDE domain has a tripartite binding site that has been the focus of drug discovery and development (Salgado-Polo and Perrakis, 2019). This tripartite site is composed of (i) a catalytic bimetallic site next to a hydrophilic shallow groove that accommodates the glycerol moiety of lipid substrates, (ii) a hydrophobic pocket that binds acyl chains, and (iii) a partially hydrophobic tunnel in a T-junction leading to the other side of the PDE domain (Hausmann et al., 2011; Nishimasu et al., 2011). Importantly, the tunnel binds LPA (Nishimasu et al., 2011) as well as steroid molecules (Keune et al., 2016), and has been proposed to serve as an ‘exit channel’ that might also modulate catalytic efficiency (Nishimasu et al., 2011; Salgado-Polo et al., 2018).

ATX interacts with cell-surface integrins (Fulkerson et al., 2011) and/or heparan sulfate proteoglycans (Houben et al., 2013), and has been speculated to directly interact with LPARs, although evidence for the latter notion is lacking (Blaho and Chun, 2018; Jethwa et al., 2016). Through interaction with the cell surface, ATX may facilitate delivery of LPA to its cognate GPCRs in a highly localized manner (Perrakis and Moolenaar, 2014). Recent studies have revealed ATX/LPA as a T-cell repellent and suggested that ATX secreted by tumor cells functions as an LPA-producing ‘chaperone’ that protects LPA from rapid degradation (Matas-Rico et al., 2021).

During the last decades, many small-molecule ATX inhibitors have been developed that show different ATX binding modes with respect to occupying the catalytic site and the hydrophobic pocket together (type I), or the pocket alone (type II), or the tunnel alone (type III) (Salgado-Polo and Perrakis, 2019). The first ATX inhibitor to enter clinical trials for idiopathic pulmonary fibrosis (IPF), the clinical candidate ziritaxestat (GLPG1690) designed by Galapagos NV (Mechelen, Belgium), is a hybrid molecule that occupies both the hydrophobic pocket and the tunnel and was defined as a type IV inhibitor. Subsequent ATX inhibitors to enter clinical trials for fibrotic diseases and tumor progression are either of type III or type IV. It remains unclear whether the ATX binding mode of the different inhibitor types determines physiological outcome and, particularly, the function of ATX’s unique tunnel therein.

Here, we address these questions by showing that only tunnel-blocking ATX inhibitors are capable of inhibiting the full spectrum of LPA-mediated signaling responses. Furthermore, we demonstrate that an intact tunnel is essential for this effect. Strikingly, ATX/LPA-mediated signaling shows a strong preference for P2Y family GPCRs (LPA_6_), which accept the LPA ligand from the lipid bilayer. Finally, we validate these findings in lung fibroblasts and in a murine model of radiation-induced pulmonary fibrosis (Bickelhaupt et al., 2017). Taken together, our results show that the ATX tunnel is essential for transport and delivery of LPA to LPARs, resulting in enhanced and selective signaling outputs. Thus, by virtue of the ATX tunnel, ATX-bound LPA functions as a ‘biased’ receptor agonist. These findings bring a new perspective in our understanding of the ATX–LPA signaling axis and highlight the clinical benefit of targeting the ATX tunnel to block LPA signaling.

## RESULTS

### Different types of ATX inhibitors modulate select signaling events

We analyzed two highly potent compounds inhibiting LPC hydrolysis: a newly developed type I inhibitor, termed Compound A (CpdA) (Fig. 1A), and the type IV inhibitor ziritaxestat (GLPG1690) (Fig. 1B). A crystal structure of CpdA (K_i_=4 nM) bound to rat ATX confirmed that the molecule is a classic competitive type I inhibitor, occupying the active site and hydrophobic pocket and forming several hydrogen bonding interactions (Fig. 1A, Fig. S1A). Ziritaxestat has a ∼25-fold lower affinity for ATX (K_i_=95 nM) and the crystal structure bound to rat ATX closely recapitulated the structure previously presented in complex with murine ATX (Desroy et al., 2017). This demonstrates that ziritaxestat is a type IV inhibitor; its binding in the ATX tunnel is mediated mostly by van der Waals interactions (Fig. 1B, Fig. S1A). As the tunnel is a secondary binding site for LPA, which can modulate LPC hydrolysis (Salgado-Polo et al., 2018), we confirm that ziritaxestat, but not CpdA, competes with LPA for tunnel occupancy (Fig. S1B).

**Figure 1.**
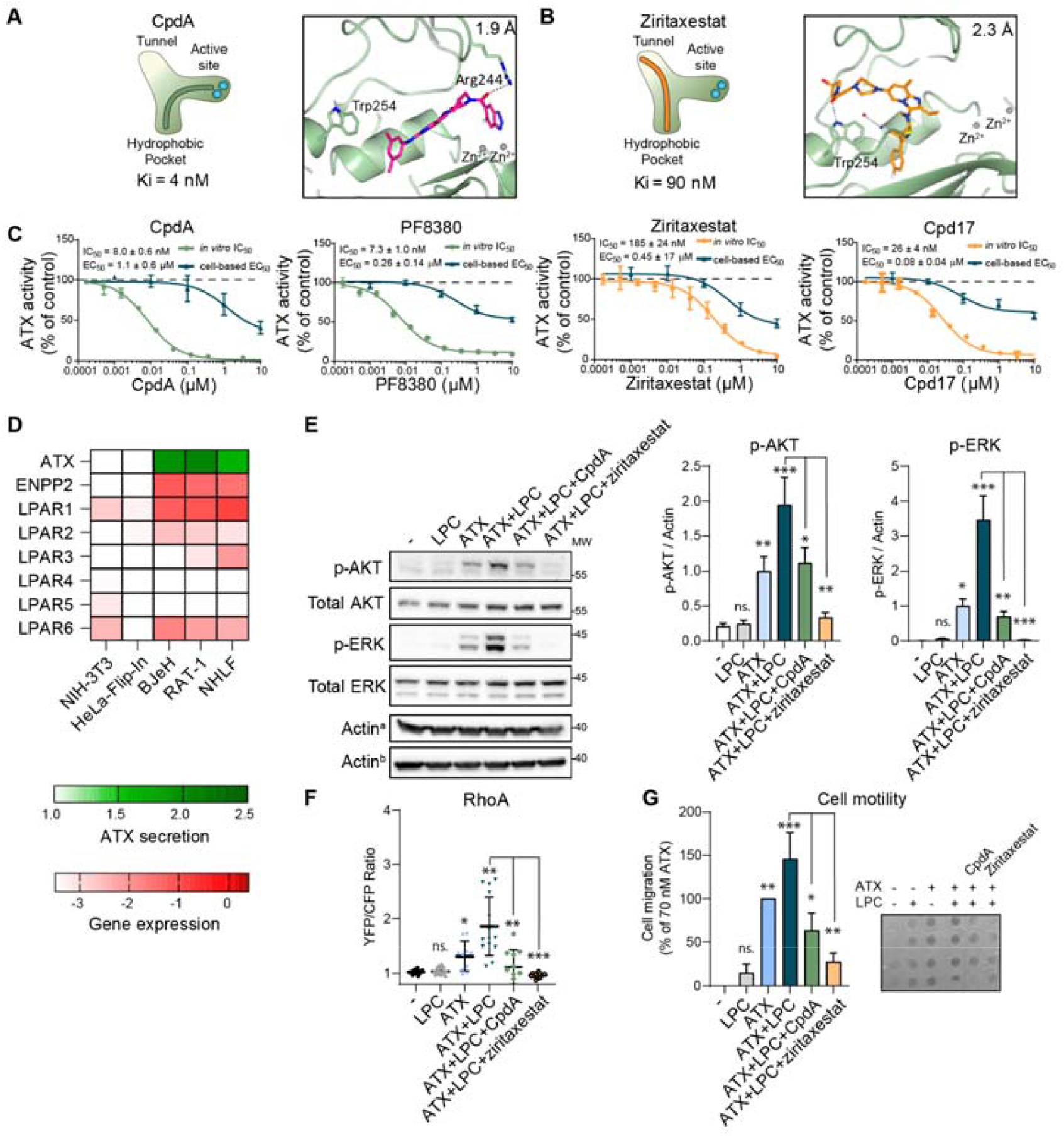
Different ATX inhibitors differentially modulate cellular signaling. (**A**,**B**) Schematic of (**A**) type I and (**B**) type IV modes of binding. Binding pose of CpdA and ziritaxestat at the ATX tripartite site from co-crystal structures (PDB ID 7Z3K, 7Z3L) are shown on the right. **(C)**ATX activity, measured by choline release from LPC(18:1), and the inhibitory effect of four different ATX inhibitors, indicated as IC_50_ values. The effect of type I (CpdA) and type IV (ziritaxestat) inhibition on ATX-mediated LPC(18:1) hydrolysis in BJeH human skin fibroblasts was examined using Western blot analysis of p-AKT, and is indicated as cell-based EC_50_ values. Data represent the mean value of triplicate measures ± SEM (error bars). **(D)**Relative qPCR expression patterns of ATX (*ENPP2*) and *LPAR1–6* in the indicated cell lines. Results are shown as a heat map where Ct values were normalized to cyclophilin and presented in logarithmic scale. ATX secretion is quantified in **Fig. S1C**. **(E)**Murine NIH-3T3 cells were treated with ATX-bound LPC (1 μM) for 5 min in the presence or absence of CpdA (5 μM) or ziritaxestat (5 μM), then analyzed by Western blotting. Left panel, representative Western blots of AKT and ERK activation (total AKT, total ERK, and actin are shown as loading controls); right panels, quantitation of p-AKT and p-ERK from three independent experiments using actin as a loading control. Data represent the average value of triplicate biological measures ± SEM (error bars). aactin from p-AKT and p-ERK blot, bactin from total AKT and total ERK blot; *p<0.05, **p<0.01, ***p<0.001; ns, not significant (unpaired t-test). **(F)**Activation of RhoA in NIH-3T3 cells, measured as YFP/CFP fluorescence ratio, upon stimulation by ATX-bound LPC (1 μM) for 10 min in the presence or absence of CpdA (5 μM) or ziritaxestat (5 μM). Data depict median ± IQR of 20 fields containing at least 10 cells. *p<0.05, **p<0.01, ***p<0.001; ns, not significant (one-way ANOVA). **(G)**Transwell migration of NIH-3T3 cells in response to albumin- or ATX-bound (20 nM) LPC (1 μM) in the lower chamber upon 4 h. ATX was allowed to hydrolyze LPC into LPA where it was not inhibited by CpdA (5 μM) or ziritaxestat (5 μM). Left panel, quantitation of stimulant-dependent cell migration; right panel, representative filter containing fixed cells. Data represent the average value of triplicate biological measures ± SEM (error bars). *p<0.05, **p<0.01, ***p<0.001; ns, not significant (unpaired t-test).

Having verified the structural binding mode for both compounds, we examined whether these distinct inhibitors behave differently in a cellular context. The effects of CpdA and ziritaxestat, in addition to PF8380 (type I inhibitor) (Gierse et al., 2010) and Cpd17 (type IV inhibitor) (Keune et al., 2017), were therefore assessed. Type I compounds were more efficient in inhibiting ATX catalytic activity *in vitro* (IC_5o_ <10 nM) than type IV compounds (IC_5o_ >25 nM) (Fig. 1C). However, type I and type IV compounds were equally proficient in inhibiting ATX signaling in cell culture (EC_50_ ∼1 μM), as measured by Western blot analysis of AKT activation in human BJeH skin fibroblasts (Fig. 1C). The lack of correlation between inhibition of catalytic efficiency *in vitro* and cellular outcome suggests that occupying the ATX tunnel could determine biological outcome.

To corroborate these findings, we examined the effect of ATX in an ATX-free extracellular environment. We examined ATX expression (by qPCR) and secretion (by Western blotting) in several cell lines (Fig. 1D, Fig. S1C). As NIH-3T3 fibroblasts were found to lack detectable ATX expression (Fig. 1D, Fig. S1C), as reported previously (Nam et al., 2000), these cells were used in subsequent experiments.

Cells were stimulated using recombinant ATX, with or without LPC substrate, alone or with ATX inhibitors CpdA or ziritaxestat. Phosphorylation of AKT and ERK was more strongly inhibited by ziritaxestat than CpdA (Fig. 1E). We noted that p-AKT or p-ERK were reduced in absolute levels (normalized to total actin), and in comparison to total AKT or ERK levels (in subsequent experiments only normalized values are reported). Moreover, ziritaxestat markedly abrogated RhoA activation, whereas the impact of CpdA was limited (Fig. 1F). ATX-induced transwell cell migration was increased by LPC(18:1) and was more efficiently reduced by ziritaxestat than by CpdA (Fig. 1G).

The feature that distinguishes ziritaxestat from CpdA is not its potency in inhibiting LPC hydrolysis, but its occupancy of the LPA-binding tunnel. This suggests that specific LPA signaling functions may be mediated through LPA binding to the ATX tunnel. We therefore examined if ATX can act as an LPA chaperone, independent of its catalytic activity.

### ATX is a dual-function protein that acts as an LPA chaperone in LPA signaling

LPA is known to activate AKT, ERK, and RhoA through distinct G protein–effector pathways. Of note, in all subsequent experiments, we used fatty acid-free albumin (a known LPA carrier) at a concentration of ∼8 μM (0.05% w/v)—much higher than the concentration of recombinant ATX (20 nM). Albumin-bound LPA efficiently activated ERK in NIH-3T3 cells, but only marginally activated AKT and RhoA (Fig. 2A,B). Addition of recombinant ATX alone activated AKT, RhoA, and, to a lesser extent, ERK (Fig. 2A,B). It therefore appears that both LPA in the presence of albumin and ATX alone (without added LPA) activate complementary pathways. Further, ATX preincubated with LPA (ATX-bound LPA) enhanced the activation of AKT, ERK, and RhoA when compared to stimulation with LPA or ATX alone (Fig. 2A,B, Fig. S2A,B). In cell migration assays, NIH-3T3 cells failed to respond to albumin-bound LPA, but did respond to ATX-bound LPA (Fig. 2C). Similar results were obtained using ATX-deficient MDA-MB-231 breast carcinoma cells (Fig. S2C). However, in ATX-secreting RAT-1 and BJeH fibroblasts, albumin-bound LPA was sufficient to activate both AKT and ERK; this activation was not enhanced by adding ATX (Fig. 2D, Fig. S2D).

**Figure 2.**
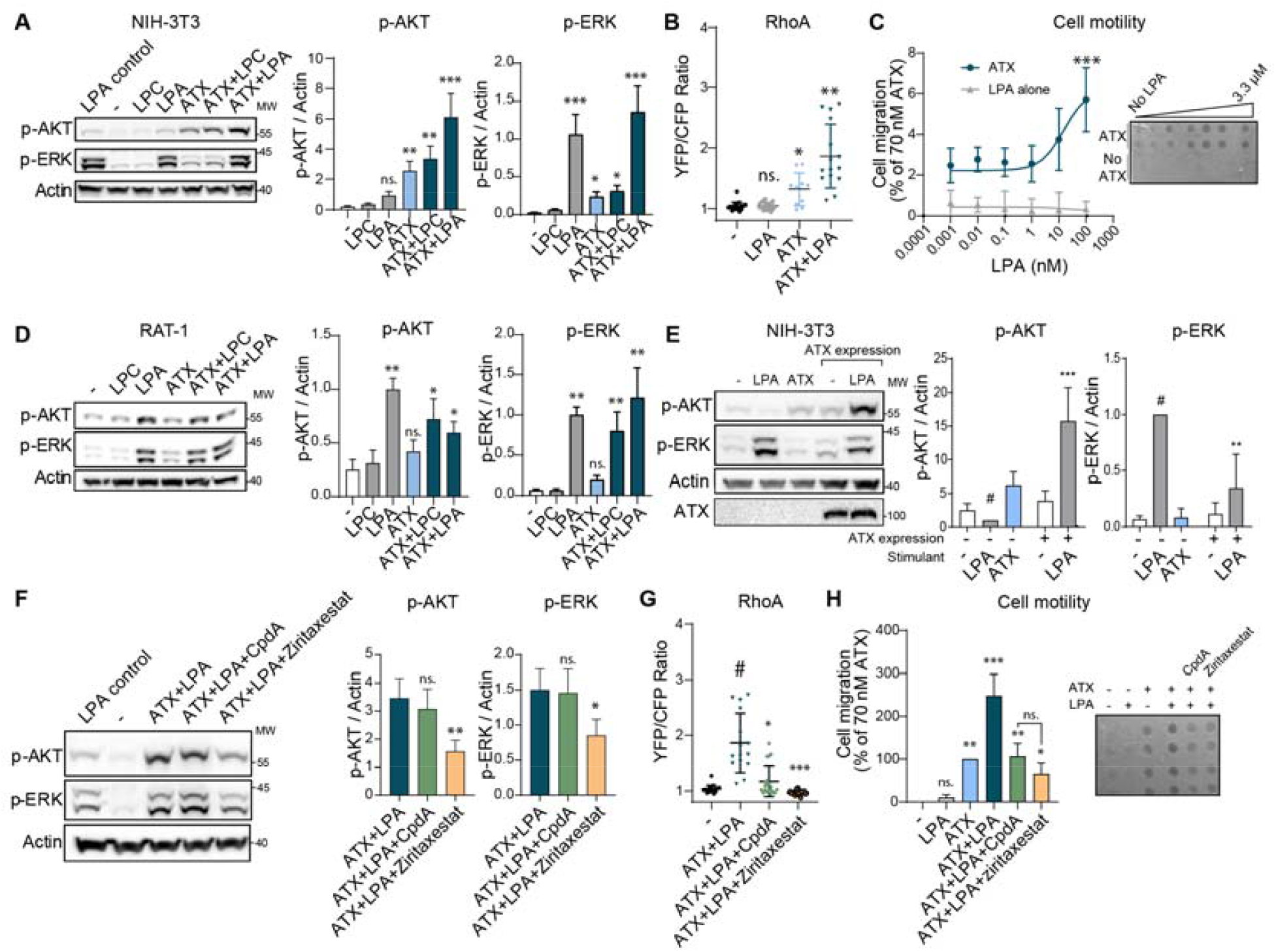
Role of ATX in LPA-mediated signaling and effect of distinct ATX inhibitors. **(A)**Role of ATX as an LPA carrier driving activation of AKT and ERK in NIH-3T3 cells. Cells were serum starved for 16 h and stimulated for 5 min with albumin-bound LPC (1 μM) or LPA (1 μM), with ATX alone (20 nM) or with ATX-bound LPC or LPA. Albumin-bound LPA was used as a control for NIH-3T3 cells after a 2.5-min stimulation and was used for normalization. Left panel, representative Western blot; right panels, quantitation of AKT and ERK activation. Three independent experiments were quantified, shown as the mean ± SEM. *p<0.05, **p<0.01, ***p<0.001; ns, not significant (one-way ANOVA). **(B)**RhoA activation in NIH-3T3 cells in response to the indicated stimulants. The response amplitude was quantitated (see **Fig. S2** for complete time course). Data depict median ± IQR of 20 fields containing at least 10 cells. *p<0.05, **p<0.01; ns, not significant (one-way ANOVA). **(C)**Boyden chamber NIH-3T3 cell migration assay performed in serum-free media, using a gradient of albumin- or ATX-bound LPA(18:1) as stimulant, where the concentrations of the carriers remained constant. Left panel, quantitation of LPA-dependent cell migration depending on the carrier; right panel, representative filter containing fixed cells. Albumin alone does not suffice to enable LPA-mediated cell migration. Data represent the average value of triplicate biological measures ± SEM (error bars). ***p<0.001 (unpaired t-test). **(D)**Role of ATX as an LPA carrier driving activation of AKT and ERK in NIH-3T3 cells. Cells were serum starved for 16 h and stimulated for 5 min with albumin-bound LPC (1 μM) or LPA (1 μM), with ATX alone (20 nM) or with ATX-bound LPC or LPA. Note the contrasting pattern of p-AKT to that of panel (**A**). Left panel, representative Western blot; right panels, quantitation of AKT and ERK activation. Quantitation of three independent experiments, shown as the mean ± SEM. *p<0.05, **p<0.01; ns, not significant (one-way ANOVA). **(E)**Need of endogenous ATX production for AKT activation in NIH-3T3 cells. Cells were serum starved and transfected with 1 μg ATX cDNA for 24 h. Left panel, representative Western blot; right panels, quantitation of AKT and ERK activation. Three independent experiments were quantified, shown as the mean ± SEM. **p<0.01, ***p<0.001 (one-way ANOVA). **(F)**The ability of CpdA and ziritaxestat to compete with LPA for binding to ATX was tested in NIH-3T3 cells. Cells were treated with ATX-bound LPA for 5 min in the presence or absence of CpdA (5 μM) or ziritaxestat (5 μM) and compared to albumin-bound LPA (control). Left panel, representative Western blot; right panels, quantitation of AKT and ERK activation. Quantitation of three independent experiments, shown as the mean ± SEM. *p<0.05, **p<0.01; ns, not significant (one-way ANOVA). **(G)**RhoA activation in NIH-3T3 cells in response to ATX-bound LPA for 10 min in the presence or absence of CpdA (5 μM) or ziritaxestat (5 μM). Data depict median ± IQR of 20 fields containing at least 10 cells. *p<0.05, ***p<0.001 (one-way ANOVA). **(H)**Transwell migration of NIH-3T3 cells in response to albumin-or ATX-bound (20 nM) LPA (1 μM) in the lower chamber. The ATX chaperone function was hampered by CpdA (5 μM) or ziritaxestat (5 μM). Left panel, quantitation of stimulant-dependent cell migration; right panel, representative filter containing fixed cells. Data represent the average value of triplicate biological measures ± SEM (error bars). *p<0.05, **p<0.01; ***p<0.001; ns, not significant (unpaired t-test).

Next, we examined if the effect of exogenously added ATX could be recapitulated by enforced ATX expression in NIH-3T3 cells. Transient ATX expression enabled LPA-induced activation of both AKT and ERK (Fig. 2E); similar to the effect observed in BJeH and RAT-1 cells expressing endogenous ATX.

This suggests that ATX is needed for specific aspects of LPA signaling. As binding of ziritaxestat to the tunnel was important for abrogating the activation of AKT, ERK, and RhoA in the context of ATX-mediated LPC hydrolysis, we asked if ziritaxestat or CpdA affect the action of ATX-bound LPA. Ziritaxestat, but not CpdA, inhibited AKT and ERK activation by ATX-bound LPA (Fig. 2F, Fig. S2E). Consistently, ziritaxestat showed a stronger effect than CpdA on RhoA activation (Fig. 2G). Both compounds affected NIH-3T3 cell migration, with ziritaxestat exerting a more pronounced effect (Fig. 2H).

Taken together, these results suggest that type IV inhibitors occupying the ATX tunnel do more than inhibit LPA production. According to this scenario, ATX is a dual-function protein that acts as an LPA-producing chaperone with functional specificity.

### LPA delivery by ATX requires a structurally intact tunnel

To decouple the catalytic activity of ATX from its chaperone signaling function, we mutated the catalytic nucleophile Thr210 to abrogate ATX lysoPLD activity, and residue Trp255 located in the LPA-binding ATX tunnel. Both mutants render ATX catalytically inactive *in vitro* (Fig. S3A).

Wildtype ATX activates AKT, but not ERK, on its own (Fig. 1E, Fig. 2A). Strikingly, but consistent with the dual-function hypothesis, catalytically inactive ATX(T210A) markedly activated AKT, although to a lesser extent than wildtype ATX, but failed to activate ERK (Fig. 3A, Fig. S3B,C). The ATX(W255A) tunnel mutant was incapable of activating AKT (Fig. 3A, Fig. S3D,E). Preincubating ATX with LPA to produce ATX-bound LPA resulted in AKT activation by both wildtype ATX and ATX(T210A), but not by ATX(W255A) (Fig. 3A, Fig. S2A, Fig. S3B–E). Notably, addition of albumin-bound LPA resulted in ERK activation, regardless of the presence of ATX or any of its mutants (Fig. 3A, Fig. S3B–E). Catalytically inactive ATX(T210A) activated RhoA, albeit to a lesser extent than wildtype ATX; by contrast, the ATX(W255A) tunnel mutant failed to activate RhoA (Fig. 3B, S3F). Additionally, in NIH-3T3 cell migration assays, both wildtype ATX and ATX(T210A), but not ATX(W255A), induced chemotactic activity when preincubated with LPA (Fig. 3C). These results indicate that catalytically inactive ATX can exert physiological activity, with a key function for the ATX tunnel.

**Figure 3.**
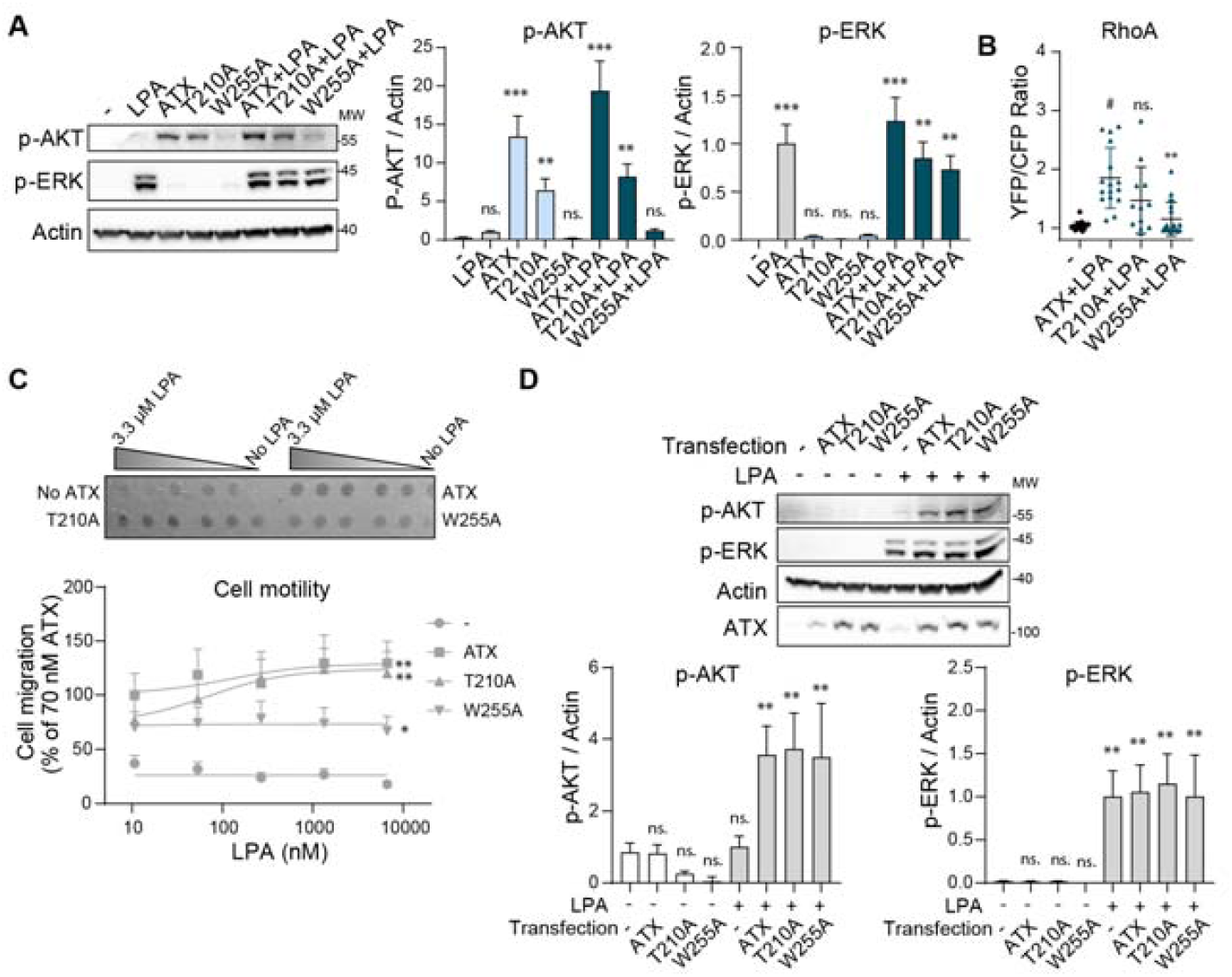
LPA delivery as a catalysis-independent event that requires an intact ATX tunnel. **(A)**AKT and ERK activation in NIH-3T3 cells in response to albumin-bound LPA (1 μM), 20 nM ATX, catalytically inactive ATX(T210A), or tunnel mutant ATX(W255A), or LPA bound to ATX, ATX(T210A), or ATX(W255A). Stimulants were preincubated for 30 min and cells were stimulated for a total of 5 min. Left panel, representative Western blot; right panels, quantitation of activated AKT and ERK. Three independent experiments were quantified, shown as the mean ± SEM. **p<0.01, ***p<0.001; ns, not significant (one-way ANOVA). **(B)**RhoA activation in NIH-3T3 cells in response to 20 nM wildtype ATX, ATX(T210A), or ATX(W255A). The response amplitude was quantitated (see **Fig. S3F** for complete time course). Data depict median ± IQR of 20 fields containing at least 10 cells. **p<0.01; ns, not significant (one-way ANOVA). **(C)**Transwell NIH-3T3 cell migration assay performed in serum-free medium, using recombinant ATX, ATX(T210A), or ATX(W255A) (20 nM) in the presence or absence of 1 μM LPA as stimulants. Upper panel, representative filter containing fixed cells; lower panel, quantitation of LPA-dependent cell migration. Data represent the average value of triplicate biological measures ± SEM (error bars). *p<0.05, **p<0.01 (unpaired t-test). **(D)**NIH-3T3 cells were serum starved and transiently transfected with 1 μg ATX, ATX(T210A), or ATX(W255A) cDNA for 24 h, then analyzed by Western blotting. Upper panel, representative Western blot of AKT and ERK activation; lower panels, quantitation of activated AKT and ERK. Quantitation of three independent experiments, shown as the mean ± SEM. **p<0.01; ns, not significant (one-way ANOVA).

We then asked if endogenously secreted ATX could recapitulate the signaling outcome triggered by recombinant protein. Transient transfection of the respective ATX variants in NIH-3T3 cells led to ATX secretion and enabled LPA-induced AKT activation (Fig. 3D). This suggests that catalytically inactive ATX can enhance LPA-induced AKT activation, leaving ERK activation unaffected. Under these conditions, ATX(W255A) and ATX(T210A) had similar effects (Fig. 3D). Although seemingly at odds with the proposed importance of an intact tunnel, this result could be explained by the observed differences in secreted ATX levels and exposure time, as ATX(T210A) and ATX(W255A) were more abundant than wildtype ATX in these studies.

### Functional selectivity of ATX: preference for P2Y-type LPA_6_ over EDG-type LPA_1_

Given that LPA_1-3_ and LPA_4-6_ are evolutionarily distinct receptors and use different ligand entry modes (Fig. 4A), we asked whether ATX/LPA shows LPAR selectivity. Since HeLa-Flp-In cells exhibited very low (or undetectable) expression of ATX and LPA_1-6_ (Fig. 1D), we used these cells to reconstitute inducible expression of EDG LPA_1_ and P2Y LPA_6_ (C-terminally HA-tagged) and analyze ATX/LPAR signaling. We confirmed that both LPA_1_ and LPA_6_ were expressed at similar levels, as shown by qPCR and Western blot analyses (Fig. S4A,B), and were localized to the plasma membrane, as shown by confocal microscopy (Fig. S4C).

**Figure 4.**
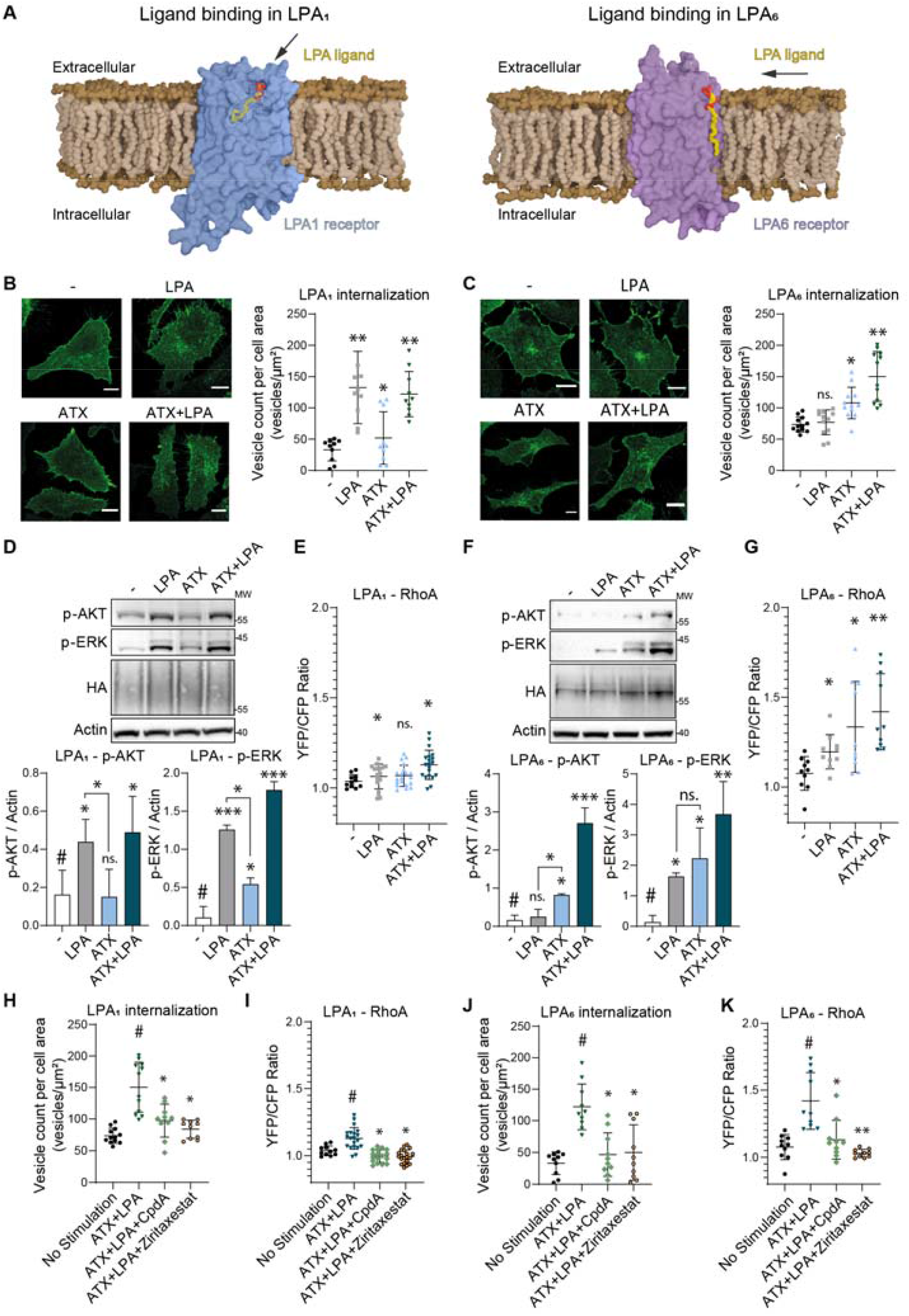
ATX shows a preference for P2Y-type LPA_6_ over EDG-type LPA_1_. (**A**) Surface representation of the differing entry modes of LPA(18:1) into the binding pockets of LPA_1_ (left) and LPA_6_ (right). UCSF ChimeraX 1.2.5 was used to generate surfaces. Membranes with bound LPARs were created and rendered using Blender 2.93.5.PDB codes. The structures were retrieved from the AlphaFold database (AF-Q92633-F1 [LPA_1_] and AF-P43657-F1 [LPA_6_] (Jumper et al., 2021; Varadi et al., 2022)). (**B, C**) Representative images and quantitation of the internalization of (**B**) LPA_1_-HA and (**C**) LPA_6_-HA in HeLa-Flp-In cells upon stimulation with albumin-bound LPA (1 μM), ATX (20 nM), or ATX-bound LPA for 15 min. Left panels, representative confocal images used for vesicle quantitation; right panel, calculation of the number of internalized vesicles. Data represent the average value ± SEM (error bars) of triplicate experiments where at least 20 fields were analyzed. *p<0.05, **p<0.01; ns, not significant (unpaired t-test). **(D)**Activation of AKT and ERK in LPA_1_-HA-expressing HeLa cells. Stimulation of LPA_1_-HA in HeLa-Flp-In cells that were starved overnight with 0.5% serum-containing medium, where receptor expression was also induced. Top panel, representative Western blot; bottom panels, quantitation of AKT and ERK activation. Quantitation of three independent experiments, shown as the mean ± SEM. *p<0.05, ***p<0.001; ns, not significant (one-way ANOVA). **(E)**RhoA activation in response to albumin-bound LPA (1 μM), ATX (20 nM), and ATX-bound LPA in HeLa-Flp-In cells (YFP/CFP fluorescence ratio) mediated by LPA_1_-HA. The response amplitude was quantitated (see **Fig. S4** for full time course). Data depict median ± IQR of 20 fields containing at least 10 cells. *p<0.05; ns, not significant (one-way ANOVA). **(F)**Activation of AKT and ERK in LPA_6_-HA-expressing HeLa cells. Stimulation of LPA_6_-HA in HeLa-Flp-In cells that were starved overnight with 0.5% serum-containing medium, where receptor expression was also induced. Top panel, representative Western blot; bottom panels, quantitation of AKT and ERK activation. Quantitation of three independent experiments, shown as the mean ± SEM. *p<0.05, **p<0.01; ***p<0.001; ns, not significant (one-way ANOVA). **(G)**RhoA activation in response to albumin-bound LPA (1 μM), ATX (20 nM), and ATX-bound LPA in HeLa-Flp-In cells (YFP/CFP fluorescence ratio) mediated by LPA_6_-HA. The response amplitude was quantitated (see **Fig. S4** for full time course). Data depict median ± IQR of 20 fields containing at least 10 cells. *p<0.05, **p<0.01 (one-way ANOVA). **(H)**Induction of the internalization of LPA_1_-HA in HeLa-Flp-In cells by ATX-bound LPA (20 nM, 1 μM) with or without CpdA (5 μM) or ziritaxestat (5 μM). Intracellular vesicle count upon a 15-min stimulation in the presence of serum-free DMEM containing 0.05% FAF-BSA. Data represent the average value ± SEM (error bars) of triplicate experiments where at least 20 fields were analyzed. *p<0.05 (unpaired t-test). **(I)**RhoA activation in LPA_1_-HA-expressing HeLa-Flp-In cells (shown as a YFP/CFP fluorescence ratio) upon stimulation with ATX-bound LPA for 10 min in the presence or absence of CpdA (5 μM) or ziritaxestat (5 μM). Data depict median ± IQR of 20 fields containing at least 10 cells. *p<0.05 (one-way ANOVA). **(J)**Induction of the internalization of LPA_6_-HA in HeLa-Flp-In cells by ATX-bound LPA (20 nM, 1 μM) with or without CpdA (5 μM) or ziritaxestat (5 μM). Intracellular vesicle count upon a 15-min stimulation in the presence of serum-free DMEM containing 0.05% FAF-BSA. Note that CpdA and ziritaxestat abolish vesicle internalization in LPA_6_-HA-expressing cells. Data represent the average value ± SEM (error bars) of triplicate experiments where at least 20 fields were analyzed. *p<0.05 (unpaired t-test). **(K)**RhoA activation in LPA_6_-HA-expressing HeLa-Flp-In cells (shown as a YFP/CFP fluorescence ratio) upon stimulation with ATX-bound LPA for 10 min in the presence or absence of CpdA (5 μM) or ziritaxestat (5 μM). Data depict median ± IQR of 20 fields containing at least 10 cells. *p<0.05; **p<0.01 (one-way ANOVA).

We used these cell lines to examine receptor activation in response to ATX, LPA, and ATX preincubated with LPA. As a readout, we used agonist–receptor internalization and quantified cytoplasmic LPAR-containing vesicles by confocal imaging. LPA_1_ was internalized in response to albumin-bound LPA and ATX-bound LPA, but to a lesser extent in response to ATX alone (Fig. 4B). By contrast, LPA_6_ did not exhibit detectable internalization upon LPA stimulation, but responded more strongly to ATX and to ATX preincubated with LPA (Fig. 4C). This suggests that the non-catalytic effects of ATX are preferentially mediated by the P2Y-type LPA receptors than by EDG-type LPA receptors.

Consistent with this, LPA_1_-expressing HeLa cells responded to albumin-bound LPA, as shown by activation of AKT, ERK, and RhoA, irrespective of ATX presence (Fig. 4D,E, Fig. S4D,E). In marked contrast, LPA_6_-expressing cells showed much stronger AKT/ERK/RhoA activation responses to ATX-bound LPA than to albumin-bound LPA (but weaker than to ATX alone) (Fig. 4F,G, Fig. S4D,E).

Finally, we examined if CpdA and ziritaxestat differentially affected LPA_1_-versus LPA_6_-mediated signaling in induced HeLa cells. We focused on the dual function of ATX, using ATX preincubated with LPA. In LPA_1_-expressing cells, both CpdA and ziritaxestat reduced the internalization of LPA_1_ and activation of RhoA (Fig. 4H,I). In LPA_6_-expressing cells, both compounds reduced LPA_6_ internalization (Fig. 4J), but ziritaxestat was a more efficient inhibitor of RhoA activation (Fig. 4K).

Collectively, these findings suggest that ATX-bound LPA signaling shows preference for P2Y LPARs. Given the specific effect of ziritaxestat and its therapeutic potential for lung fibrosis, we aimed to corroborate the above mechanism in primary human lung fibroblasts.

### ATX-mediated LPA delivery in human lung fibroblasts

Normal human lung fibroblasts (NHLFs) express and secrete ATX and express high levels of EDG receptors LPA_1_ and LPA_3_, and P2Y receptor LPA_6_ (Fig. 1D).

Treatment of NHLFs with albumin-bound LPA or LPC resulted in weak activation of AKT; albumin-bound LPA also activated ERK (Fig. 5A, S5A). However, ATX alone, ATX plus LPC, and ATX-bound LPA activated both AKT and ERK. This is in agreement with the results from NIH-3T3 cells (Fig. 2A). Stimulation with catalytically inactive ATX(T210A) resulted in AKT activation, whereas stimulation with the tunnel mutant ATX(W255A) failed to do so (Fig. S5B). CpdA did not inhibit AKT or ERK activation (Fig. 5B), and only affected the RhoA response to ATX-bound LPA to a low extent (Fig. 5C, Fig. S5C). By contrast, tunnel-binding ziritaxestat abrogated AKT and RhoA activation, confirming that our results extend to early-passage NHLFs. As all cell-based assays confirmed that ATX is a dual-function protein, and that ziritaxestat (but not CpdA) inhibits its activity as an LPA chaperone with functional specificity (Fig. 5D), we then examined the relative efficacy of ziritaxestat and CpdA *in vivo*, in a murine model of pulmonary fibrosis.

**Figure 5.**
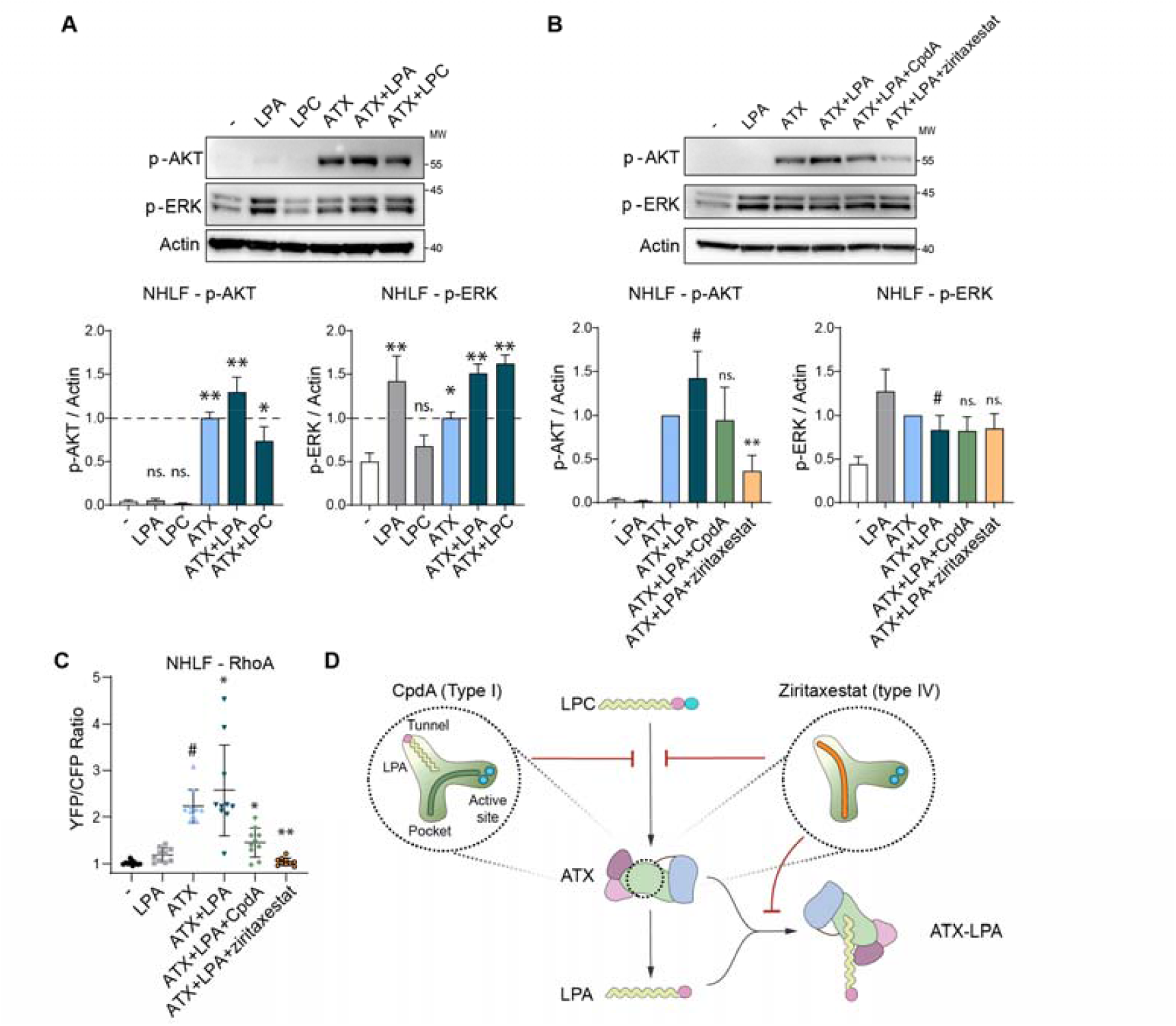
ATX-mediated LPA delivery and signaling in human lung fibroblasts. **(A)**Stimulation of NHLFs with LPA or LPC in the presence or absence of ATX. Top panel, representative Western blot; bottom panels, quantitation of AKT and ERK activation. Quantitation of three independent experiments, representing the average value of triplicate biological measures ± SEM (error bars). *p<0.05, **p<0.01, ns, not significant (unpaired t-test). **(B)**Stimulation of NHLFs with LPA, ATX, or ATX-LPA preincubated with or without CpdA (5 μM) or ziritaxestat (5 μM). Top panel, representative Western blot; bottom panels, quantitation of AKT and ERK activation. Quantitation of three independent experiments, representing the average value of triplicate biological measures ± SEM (error bars). **p<0.01, ns, not significant (unpaired t-test). **(C)**RhoA activation in response to albumin-bound LPA, ATX, or ATX-bound LPA for 10 min in the presence or absence of CpdA (5 μM) or ziritaxestat (5 μM). The response amplitude was quantitated (see **Fig. S5** for full time course). Data depict median ± IQR of 20 fields containing at least 10 cells. *p<0.05, **p<0.01 (one-way ANOVA). **(D)**Schematic representation of secreted ATX with its tripartite site, LPA binding in the ATX tunnel, and the binding modes of ATX inhibitors CpdA and ziritaxestat. LPA_1_ accepts LPA from the water phase, whereas LPA_6_ accepts LPA from the lipid bilayer (**Fig. 4A**), with ATX serving as an LPA-producing chaperone.

### Tunnel-blocking ziritaxestat, but not CpdA, reverses pulmonary fibrosis in mice

Ziritaxestat and CpdA were evaluated in a model of radiation-induced pulmonary fibrosis, a serious adverse effect of radiotherapy for patients with lung cancer with few treatment options (Ding et al., 2013; Jin et al., 2020), and relevant for progressive lung fibrosis in a therapeutic setting (Fig. 6A) (Jin et al., 2020). In our mouse model, both compounds were evaluated in two separate experiments, where the doses were selected based on the compounds’ potency in rat plasma LPA assays (Fig. 6B). Drug exposure was measured at steady state in rat plasma LPA assays, which confirmed the similar target coverage and respective IC_50_ values of the compounds (541 nM for ziritaxestat and 13.6 nM for CpdA) (Fig. 6B, Table S1). When assayed above their IC_50_ values, CpdA was more efficient than ziritaxestat in inhibiting LPA production (Fig. 6C).

**Figure 6.**
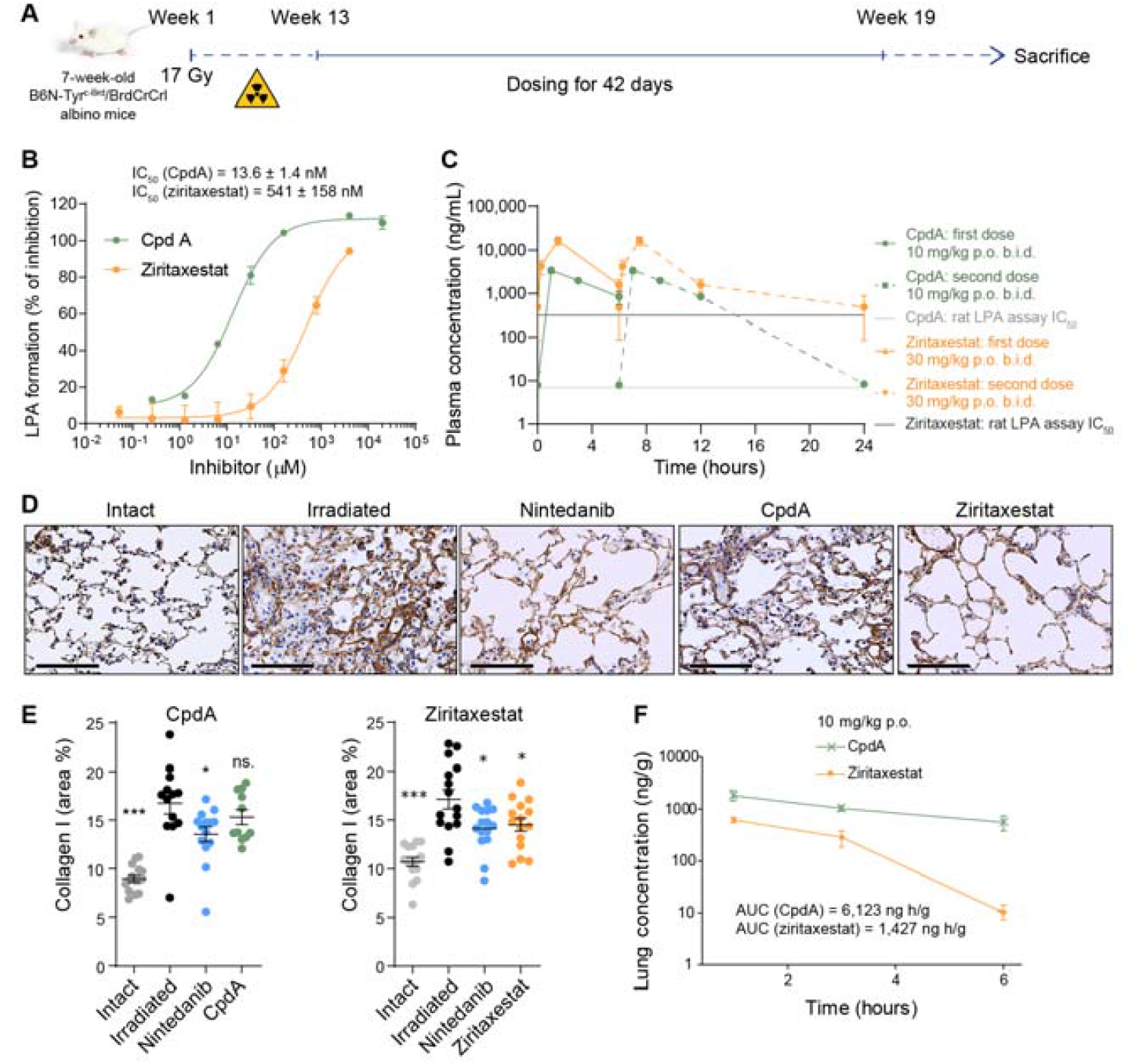
Ziritaxestat, but not CpdA, reverses pulmonary fibrosis *in vivo*. **(A)**Radiation-induced fibrosis model. Irradiated lung was treated with CpdA (10 mg/kg b.i.d.), ziritaxestat (30 mg/kg b.i.d.), or nintedanib, a clinically approved drug for the treatment of IPF (positive control; 60 mg/kg q.d.). **(B)**Percentage of reduction in 18:2 LPA formation from *ex vivo* rat plasma samples upon incubation with CpdA or ziritaxestat, as determined by LC-MS. The percentage of inhibition was calculated by normalization of data to LPA levels after a 2-h incubation at 37 °C in the absence of the compounds. The resulting IC_50_ values (± SEM) are shown. **(C)**Mean (± SEM) steady-state plasma concentrations of CpdA and ziritaxestat after a p.o. administration at 10 mg/kg b.i.d. and 30 mg/kg b.i.d., respectively, shown on logarithmic scale. (**D**,**E**) Analysis of collagen I levels in the murine model of radiation-induced fibrosis. (**D**) Representative images of collagen I staining. (**E**) Surface area quantitation of collagen I levels of CpdA- or ziritaxestat-treated mice after 19 weeks. *p<0.05, ***p<0.001; ns, not significant (one-way ANOVA with Dunnett’s post-test statistical analysis). (**F**) Lung tissue exposure of CpdA and ziritaxestat after a single p.o. administration at 10 mg/kg.

As fibrotic areas are characterized by increased secretion of extracellular matrix proteins, including type I and III collagen, we quantitated the effect of CpdA and ziritaxestat on reducing pulmonary collagen I levels. Irradiation induced a significant increase in collagen I levels after 19 weeks compared with the non-treated control, which was reduced in both experiments by treatment with nintedanib, a clinically approved drug for the treatment of IPF (Fig. 6D,E). While CpdA treatment did not decrease collagen I levels significantly, ziritaxestat treatment markedly reduced collagen I secretion (Fig. 6D,E). Pharmacokinetic analysis of lung exposure after single oral administration of both compounds (at 10 mg/kg) showed that the values for area under the concentration–time curve from 0 to 6 hours (AUC_0-6h_) were five-fold higher for CpdA than for ziritaxestat (6,123 ng.h/g vs 1,427 ng.h/g) (Fig. 6F). As such, the lack of CpdA therapeutic activity was not due to lower pulmonary exposure.

In summary, targeting the ATX tunnel and the related non-catalytic ATX activities is important both *in vitro* and *in vivo*, highlighting the therapeutic potential of ATX inhibitors that block the tunnel and its chaperone function.

## DISCUSSION

Following its implication in tumor progression and pulmonary fibrosis, ATX has attracted considerable interest as a drug target from pharmaceutical companies and academic researchers. Structural studies demonstrated that ATX inhibitors exhibit different binding modes. ATX inhibitors entering clinical trials in patients with pulmonary fibrosis or cancer include ziritaxestat, which progressed to phase 3. BBT-877 (Bridge Biotherapeutics Inc., Pangyo, South Korea), IOA-289 (iOnctura SA, Geneva, Switzerland), and cudetaxestat (Blade Therapeutics Inc., South San Francisco, USA) are now in phase 1/2 trials and are also type IV (or perhaps, for some, type III) inhibitors, displaying the common characteristic of occupying the ATX tunnel. None of the numerous potent type I orthosteric ATX inhibitors occupy the tunnel (iOnctura, 2021; Jia et al., 2021; Mulholland et al., 2020; Salgado-Polo and Perrakis, 2019), and none have entered clinical trials, to the best of our knowledge. These data raise the intriguing questions of whether and how the specific drug binding mode determines the physiological outcome of ATX/LPA signaling, both *in vitro* and *in vivo*.

Here we have addressed this question by examining how type I and type IV ATX inhibitors affect specific LPAR-mediated signaling events.

We find that type IV compounds are more efficient in inhibiting AKT activation, RhoA activation, cell migration, and receptor internalization. Strikingly, our data suggest that ATX is not only needed for LPA production, but that it can also act as an LPA chaperone. It is this specific chaperone activity that can be targeted by type III or type IV compounds, which occupy the ATX tunnel. We corroborate these findings in fibroblast cell lines, LPA_1_- and LPA_6_-expressing HeLa cells, NHLFs, and in a murine model of radiation-induced pulmonary fibrosis.

Our findings support the view that ATX acts as a dual-function protein that can activate signaling pathways by virtue of its unique tunnel, independent of its catalytic activity. The LPA chaperone role has unique characteristics, but also analogies to other lipid chaperones. Albumin is the primary and most abundant LPA chaperone, showing high LPA binding affinity. In our experiments, the albumin concentration was ∼8 μM. The chaperone function of ATX was prominent at as low as 20 nM, in the presence of albumin as a ‘competitor’ for LPA recruitment. Analogies with the S1P lysolipid chaperone are thus apparent. Serum albumin binds S1P, but with lower affinity than HDL-associated apolipoprotein M (ApoM), which serves as the major circulatory S1P chaperone (Cartier and Hla, 2019; Christoffersen et al., 2011). It has been shown that the ApoM/S1P complex evokes S1PR1-mediated biological responses in a selective manner, distinct from those evoked by albumin-bound S1P. ApoM/S1P thus serves as a ‘biased agonist’ for S1PR1. Unlike ApoM-bound S1P, albumin-bound S1P is short-lived in plasma and activates S1PR1 in a manner distinct from that of ApoM-S1P (Galvani et al., 2015). The present results for LPA and ATX point to a similar mechanism, with the distinct peculiarity of the dual enzymatic and chaperone role of ATX.

The additional role of ATX mediating LPA delivery involves specific LPA binding to the ATX tunnel, which is in agreement with a previous hypothesis (Nishimasu et al., 2011) and with biochemical evidence and molecular dynamics simulations indicating that the tunnel acts as a secondary binding site (Salgado-Polo et al., 2018). Moreover, LPA binds to and is released from the orthosteric site with t_1/2_ ≈ 0.1 s and from the tunnel with t_1/2_ ≈ 0.01 s, suggesting that LPA is more readily transferred from the tunnel for delivery to its cognate receptors (Tokumura et al., 2002). Thus, based on the available evidence, both the tunnel and the hydrophobic LPC/LPA-binding pocket can deliver receptor-active LPA. However, our experiments suggest that an intact tunnel is necessary for this to occur.

A further novel finding of this study is that the chaperone function of ATX shows high specificity for LPA_6_ over LPA_1_, at least in the context of our reconstituted system, which is supported by receptor expression patterns in tested cell lines. This observation poses the question of whether this selectivity is related to how EDG and P2Y LPARs recognize and accept LPA ligands. The crystal structures of LPA_1_ (Chrencik et al., 2015) and LPA_6_ (Taniguchi et al., 2017) have suggested two distinct entry modes for LPA: EDG family receptors bind LPA in a well-defined pocket facing the extracellular milieu, while for P2Y family receptors, a pocket is open towards the lateral membrane. It therefore appears likely that ATX binding to the plasma membrane enables efficient LPA transfer to LPA_6_ via the lipid bilayer. Another possible explanation is an indirect interaction between LPA-bound ATX and P2Y LPARs, mediated by cell-surface integrins (Fulkerson et al., 2011). Such a pathway could involve β-arrestin-mediated ‘biased’ signaling, as exemplified by the ApoM-S1PR1 signaling axis (Cartier and Hla, 2019; Christoffersen et al., 2011; Galvani et al., 2015).

Nonetheless, our results suggest that EDG LPARs are preferentially activated by albumin-bound LPA (Fleming et al., 2016), whereas P2Y LPARs prefer ATX-bound LPA. Interestingly, during early embryogenesis, ATX signals through P2Y LPA_4_ and LPA_6_, with no apparent role for EDG-type LPARs (van Meeteren et al., 2006; Yasuda et al., 2019). P2Y receptors LPA_5_ and LPA_6_ also play unique roles in anti-tumor immunity, regulating T cell receptor function and mediating ATX-induced repulsion of tumor-infiltrating T cells, respectively (Matas-Rico et al., 2021; Mathew et al., 2019).

The relevance of EDG family LPA_1_ in IPF and systemic sclerosis has been established using *Lpar1* knockout mice; these animals are protected from fibrosis in models of lung and dermal fibrosis. ATX expression is increased in the fibrotic skin of patients with systemic sclerosis and in a bleomycin mouse model of systemic sclerosis. Increased ATX and LPA levels have also been found in lung tissue and bronchoalveolar lavage fluid, respectively, of patients with IPF. Ziritaxestat has been trialed as a treatment for IPF and systemic sclerosis; however, phase 3 trials in patients with IPF were discontinued because the benefit-risk profile of the drug no longer supported continuation. Our present data support the view that the positive physiological outcome of ziritaxestat treatment is attributable to its ATX binding mode, occupying the ATX tunnel and thereby inhibiting specific signaling events including RhoA-mediated cytoskeletal reorganization. At first sight, this appears at odds with the role of EDG LPARs in IPF, as ziritaxestat shows functional preference for the prototypic P2Y LPAR, LPA_6_. However, ziritaxestat also affects LPA_1_-mediated signaling, which will contribute to its therapeutic effect. LPA_6_ is also present in lung fibroblasts (Uhlén et al., 2015). Additionally, the therapeutic effect of LPA receptor antagonists in fibrotic diseases, but also in cancer, must be viewed in the context of an interplay with the immune system; LPA_6_ is most highly expressed in immune cells (Matas-Rico et al., 2021).

Future structural and functional studies should help to elucidate the mechanism of ATX-mediated transfer of bound LPA to the membrane outer leaflet. The involvement of biased signaling, or the role of other ATX co-receptors such as integrins, could be explored. This might also advance our broader understanding of how lipid mediators, such as LPA, regulate biological functions in a receptor-selective manner. New therapeutic opportunities could be suggested, in connection with receptor expression patterns. In a more general context, this study highlights how structural and mechanistic insights may translate into the (pre)clinical efficacy of inhibitors blocking the ATX tunnel. Several inhibitors that occupy the ATX tunnel are currently in clinical trials; this work provides a rationale for their mode of action, targeting both the enzymatic and the chaperone function of ATX.

## Supporting information

Detailed Experimental Procedures

## AUTHOR CONTRIBUTIONS

Conceptualization, F.S.-P., A.P.; Formal analysis, F.S.-P., R.B.; Investigation, F.S.-P., R.B.; *In vivo* studies, F.M., C.J., L.W., P.F., B.H.; Writing – original draft, F.S.-P.; Writing – review & editing, F.S-P., W.H.M., B.H., A.P.; Supervision B.H., A.P.

## ACKNOWLEDGMENTS

High resolution mass spectral data were obtained at the EPSRC Mass Spectrometry facility at the University of Swansea, UK. X-ray diffraction data were collected at the European Synchrotron Radiation Facility (ESRF) and the Swiss Light Source (SLS). This work benefited from access to the NKI Protein Facility at Instruct-NL, an Instruct-ERIC center. We thank Elisa Matas-Rico and Willem-Jan Keune for help in establishing cell-based experiments, and Kees Jalink for discussions, ideas, and signaling reagents. This study was partially funded by Galapagos NV (Mechelen, Belgium). Editorial support was provided by Iain Haslam, PhD (Aspire Scientific, Bollington, UK), funded by Galapagos NV.

## STAR METHODS

### Resource availability

#### Lead contact

Further information and requests for resources and reagents should be directed and will be fulfilled by the lead contact, Anastassis Perrakis.

#### Materials availability

This study did not generate new unique reagents.

#### Data and code availability

The co-crystal structures of ATX bound to CpdA or ziritaxestat were deposited at the Protein Data Bank under the codes 7Z3K and 7Z3L, respectively.

- Crystallographic data have been deposited at the Protein Data Bank and are publicly available as of the date of publication. Accession numbers are listed in the key resources table.
- This paper does not report original code.
- Any additional information required to reanalyze the data reported in this paper is available from the lead contact upon request.

### Experimental model and subject details

#### Radiation-induced fibrosis mice model

Procedure and drug-naïve seven-week-old female B6 albino (B6N-Tyrc-Brd/BrdCrCrl) mice (16-18 grams) from Charles River (France) were delivered at the Plateforme de Radiothérapie Expérimentale in Institut Curie (Orsay, France). After a 7-day acclimatization period, they were identified using the micro-tattoo Aramis system. Mice were anesthetized with isoflurane, placed in holders, and irradiated with a single 17-Gy dose in the thorax area (Week 1) (Favaudon et al., 2019). The week after irradiation, mice were shipped to Galapagos Animal Facilities (Romainville, France), where they were housed 10 per cage after individual identification using micro-chip (Biolog-id, Boulogne-Billancourt, France). They were maintained at 22 °C on a 12-h light/dark cycle (07:00–19:00); food and water were provided *ad libitum*. The study was performed according to the Animal Institutional Care and Use Committee of Galapagos, ethical guidelines of animal experimentation, and the guidelines for welfare of animals in experimentation. At Week 13, fur was removed on the thorax of the anaesthetized mice (using a shaving razor then hair removing cream). *In vivo* imaging using a proprietary collagen near-infrared fluorescent probe was used to exclude mice presenting liver fibrosis, and to allocate them to each experimental group according to their lung fibrosis level. Treatment with ziritaxestat (30 mg/kg b.i.d) or CpdA (10 mg/kg b.i.d) was assayed by oral route, in comparison to nintedanib (60 mg/kg q.d). Treatment was initiated 13 weeks after irradiation and lasted for 6 weeks. On Week 17, 4 weeks after initiation of dosing, steady-state pharmacokinetics were assessed, with sampling at four time points: 0, 1, 3, and 6 h (assuming 24 h is t=0 h). Plasma was prepared and kept at −20 °C until quantification using liquid chromatography– mass spectrometry. At Week 19, mice were killed by elongation. Entire lungs were removed, rinsed with PBS, and weighed. Lobes of the lungs were separated and fixed in formaldehyde for 48 h before paraffin embedding. For every lung, 4-µm thick sections were immunostained with anti-collagen I antibody. The immunohistochemistry samples were processed automatically using the immunostainer BOND-RX (Leica). The stained sections were scanned (NanoZoomer, Hamamatsu Photonics K.K.) before quantification by image analysis (CaloPix, Tribun Health). Data were expressed as percentage of collagen I area per area of lung tissue devoid of the main constitutive collagen in vessels and bronchi.

### Method details

#### Protein production and crystallization

ATX was overexpressed and purified as described previously (Hausmann et al., 2010). For the crystallization studies, ATX was incubated with each screened compound at a 1:10 (protein:compound) ratio for at least 30 min. Crystals were grown for a least 7 days in a 24-well optimization screen (18−20% PEG 3350, 0.1−0.4 M NaSCN, and 0.1−0.4 M NH_4_I). In all cases, the best diffracting crystals were obtained at RT by mixing 1 μL of the protein:compound solution and 1 μL of reservoir solution. All crystals were vitrified in cryoprotectant, which consisted of reservoir solution with the addition of 20% (v/v) glycerol. Other solvent:protein ratios tested per condition were 1:2 and 2:1.

#### Crystallographic data and methods

The X-ray diffraction data for the ATX−inhibitor complexes were collected at SLS on beamline PXIII (Kabsch, 2010) at 100 K, and were recorded on a PILATUS 2M-F detector to high resolution. All data were processed and integrated with XDS (Kabsch, 2010). CpdA and ziritaxestat were processed on site, using the SLS automated processing pipeline, and scaled with AIMLESS (Evans, 2011). The structures were determined by molecular replacement using MOLREP (Vagin and Teplyakov, 2010), with the structure of ATX (PDB 2XR9) as the search model. Model building and subsequent refinement were performed iteratively with COOT (Emsley et al., 2010), REFMAC5 (Murshudov et al., 2011), and PDB_REDO (Joosten et al., 2014). Structure validation was carried out by MolProbity (Chen et al., 2010). The structure models and experimental diffraction data of CpdA and ziritaxestat were deposited at the Protein Data Bank under the codes 7Z3K and 7Z3L, respectively (Supplemental Table 2).

**Supplemental Table 2.** Crystallographic details.

| Data Collection | CpdA (7Z3K) | GLPG1690 (7Z3L) |
| --- | --- | --- |
| Wavelength (Å) | 1.0000 | 1.0000 |
| Resolution (Å) | 2.00 | 2.40 |
| Space group | P1 | P12<sub>1</sub>1 |
| Unit cell a b c (Å), $\beta$ (deg) | 53.7 62.5 63.6 104.2 98.6<br>93.2 | 62.8 88.7 77.0 90.0 102.8<br>90.0 |
| Data quality statistics<sup>a</sup> |  |  |
| CC<sub>1/2</sub> | 0.985 (0.437) | 0.996 (0.766) |
| R<sub>merge</sub> | 0.137 (0.798) | 0.081 (0.752) |
| $\langle I/\sigma \rangle$ | 5.6 (1.3) | 8.2 (1.6) |
| Completeness (%) | 98.0 (96.9) | 99.8 (99.5) |
| Redundancy | 3.1 (3.2) | 3.4 (3.5) |
| Refinement |  |  |
| No. atoms | 7130 | 6676 |
| protein | 6404 | 6348 |
| ligand/metal/glycan | 173 | 204 |
| water/iodine | 553 | 124 |
| R<sub>work</sub>/R<sub>free</sub> (%) | 20.2/24.2 | 24.4/25.9 |
| Validation |  |  |
| rmsd/rmsZ bond lengths (Å) | 0.0027/ 0.196 | 0.0017/0.128 |
| rmsd/rmsZ bond angles (deg) | 1.218/ 0.719 | 1.194/0.690 |
| Ramachandran preferred/outliers (%) | 95.68/0.00 | 94.24/0.26 |
| Ramachandran Z score | -0.86 | -2.21 |
| Rotamers preferred (%) | 95.68 | 93.81 |
| MolProbity/Clash score (%ile) | 96/99 | 100/100 |

<sup>a</sup>High resolution shell in parentheses.

#### AKT and ERK phosphorylation

A total of 100,000 BJeH or 50,000 HeLa cells were seeded in 12-well tissue culture plates and allowed to grow for 24 h in DMEM (Gibco, Life Technologies) containing 10% FCS (Thermo Scientific, USA) and 100 U mL−1 streptomycin/penicillin (Gibco, Life Technologies). In the case of NIH-3T3, 300,000 cells were plated on 6-well plates. Next, the cells were washed twice with PBS and serum starved overnight. ATX (20 nM) was incubated with inhibitors for 30 min in serum-free medium containing 0.05% (w/v) fatty acid-free BSA (total volume 1 mL). Medium from the 12-well plates was removed and replaced with 1 mL of the ATX–inhibitor mixture. Cells were stimulated for 15 min, after which medium was removed and the plates immediately frozen on dry ice and stored at −80 °C. For Western blotting, cells were washed with cold PBS, lysed in RIPA buffer, supplemented with protease inhibitors (20 mM NaF and 1 mM orthovanadate; Pierce), and centrifuged. Protein concentration was measured using a BCA protein assay kit (Pierce), after which LDS sample buffer (NuPAGE, Invitrogen) and 1 mM dithiothreitol (DTT) were added to the lysate. SDS-PAGE pre-cast gradient gels (4–12% Nu-Page Bis-Tris, Invitrogen) were loaded with 20 μg of total protein, followed by transfer to nitrocellulose membrane. Non-specific protein binding was blocked by 5% BSA in PBS-Tween (0.1%); primary antibodies (D9L: phospho-AKT (Ser473), 1:1,000; 4370S: phospho-ERK1/2 (Thr202/Tyr204), 1:1,000; Cell Signaling Technology) were incubated overnight at 4 °C in PBS-Tween with 5% BSA containing 0.1% NaN_3_. Horseradish peroxidase-conjugated secondary antibodies (DAKO, Glostrup, Denmark) were incubated for 1 h at room temperature (RT; 293 K) in PBS-Tween with 2.5% BSA and developed using ECL Western blot reagent.

#### Rho GTPase biosensor

A FRET pair consisting of RhoA-Cerulean3 and PKN fused to circularly permutated Venus was used (Kedziora et al., 2016). The HR1 region of PKN was used as the effector domain for activated RhoA. Experiments were performed in phenol red-lacking DMEM medium at 37 °C. Cells were allowed to adhere overnight on uncoated coverslips, after which they were serum starved and transfected with the biosensor for 24 h. Next, the coverslips were placed on a thermostat-controlled (37 °C) inverted Nikon Diaphot microscope and excited at 425 nm. Donor and acceptor emission were detected simultaneously with two photomultipliers, using a 505-nm beam splitter and optical filters: 470 ± 20 nm (CFP channel) and 530 ± 25 nm (YFP channel). The emission data were analyzed using the Fiji software and normalized to control cells. At least three independent experiments were analyzed for every condition (20 fields of view/condition, 3–5 cells/field of view, >50 cells/condition). FRET was expressed as the ratio between acceptor and donor signals, set at 1 at the onset of the experiment.

#### Surface biotin labeling

LPA_1_-HA-expressing HeLa-Flp-In cells were induced with 1 μg/mL doxycycline for 48 h; first, they were grown on 6-well plates for 24 h, next grown overnight in 0.5% FCS-containing DMEM, and lastly serum starved for 1 h. Between each of the previous steps, cells were washed twice with PBS. For LPA_1_-HA internalization, cells were stimulated with the indicated reagents in serum-free DMEM containing 0.1% fatty acid-free BSA for 15 min. Next, the cells were transferred to ice, washed in ice-cold PBS, and surface labeled with 0.2 mg/mL Sulfo-NHS-SS-Biotin (Thermo Scientific) for 30 minutes. Biotin-labeled LPA_1_-HA was detected using streptavidin beads (Pierce) and conjugated anti-HA antibody (3F10, 1:1,000; Roche Diagnostics).

#### Production of LPAR-expressing HeLa-Flp-In cells

cDNA containing human *LPAR1, LPAR2, LPAR3*, and *LPAR6* was amplified by PCR to remove its stop codon and add the restriction sites for BamHI and XhoI (or NotI for *LPAR6*), after which it was subcloned in an in-house produced pDNA5.1-HA vector. Codon-optimized gene blocks for *LPAR4* and *LPAR5* were ordered to facilitate amplification and cloning strategies, which included the addition of restriction sites for BamHI and XhoI (or NotI for *LPAR5*). For plasma membrane localization, the signal peptide of neuronal acetylcholine receptor subunit alpha-7 was added at the N-terminus of P2Y LPARs; 0.2 μg of the resulting vectors and 1.8 μg pOG44 Flp-Recombinase Expression Vector (Invitrogen) were incubated with 6 μL Fugene6 (Invitrogen) in 200 μL OptiMEM (Gibco) for 30 minutes, after which the mix was added to previously produced HeLa-Flp-In cells. Cell medium was refreshed 24 h later, and selection with 1 μg/mL puromycin was started and maintained with resistant cells.

#### Real-time quantitative PCR

Cells were allowed to grow to almost complete confluency on 10-cm dishes in 10% FCS-containing DMEM. Total RNA was extracted using the GeneJET purification kit (Fermentas). cDNA was synthesized by reverse transcription from 5 μg RNA using First Strand cDNA Synthesis Kit (Thermo Scientific). RTqPCR was performed on a 7500 Fast System (Applied Biosystems) as follows: 95 °C for 2 min, 95 °C for 0 min, 60 cycles at 95 °C for 15 s, followed by 60 °C for 1 min for annealing and extension. The final reaction mixtures (12 μL) consisted of diluted cDNA, 16SYBR Green Supermix (Applied Biosystems), 200 nM forward primer, and 200 nM reverse primer. Reactions were performed in 384-well plates, with three independent biological replicas. As a negative control, the cDNA was replaced by milliQ water. Cyclophilin was used as reference gene. Each sample was analyzed in triplicate and the normalized expression (NE) data were calculated using the equation NE= 2 –(Ct target – Ct reference).

#### Cell migration assays

Cell migration was measured using 48-well chemotaxis chambers (Neuro Probe, Inc.) equipped with 8 mm-pore polycarbonate membranes, coated with fibronectin (20 mg/mL). Cells (4.8×106/mL) were added to the upper chamber. Fatty acid-free BSA (0.5 mg/mL) was used as a lysophospholipid carrier. For NIH-3T3 cells, migration was allowed for 4 h at 37 °C in humidified air containing 5% CO_2_. Migrated cells were fixed in Diff-Quik Fix and stained using Diff-Quik II. Migration was quantified by color intensity measurements using Adobe Photoshop.

#### Detection of rat LPA plasma levels

Whole blood was collected from rats in sodium heparin tubes. The samples were centrifuged at 3,000 rpm for 15 min at 4 °C to separate the plasma, which was stored at −80 °C. CpdA and ziritaxestat were serially diluted in DMSO and 0.5 µL of the dilutions dispensed into 96-well plates placed on ice; plasma was thawed on ice. Next, 49.5 µL of plasma were added to wells containing 0.5 µL of ziritaxestat or CpdA (1% final DMSO concentration). The plates were covered with a polypropylene lid and incubated for 2 h at 37 °C in humidified air containing 5% CO_2_, with gentle shaking (except for the control samples, which were stored at −20 °C). The control samples were thawed on ice, then transferred to the incubated plates before liquid chromatography–mass spectrometry analysis. For the analysis, 10 µL of plasma proteins from the incubated plates were precipitated with an excess of methanol containing the internal standard 17:0 LPA (Cat# 857127, Avanti Polar Lipids, Inc.). The samples were centrifuged and the supernatants injected onto a C18 column. Analytes were eluted off the column under isocratic conditions. An API5500 QTRAP mass spectrometer (ABSciex™) was used for the detection of 18:2 LPA. Relative quantities were calculated based on the peak area.

#### Organic chemistry methods

##### Structure of GLPG1690 and CpdA

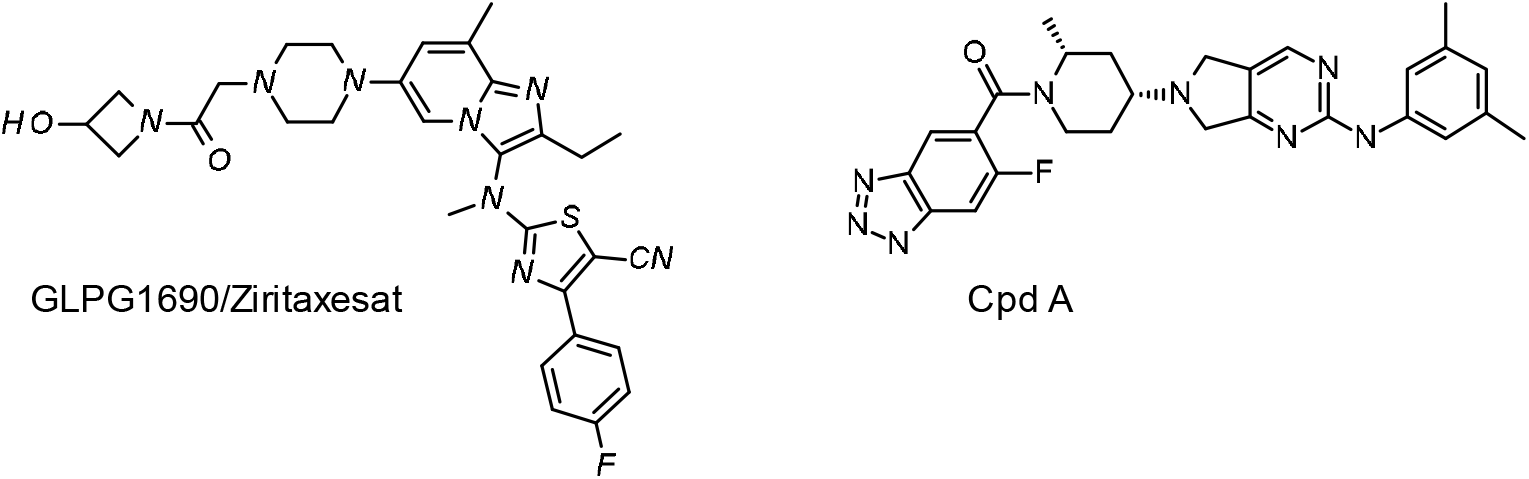

GLPG1690/ziritaxestat synthesis was published in (Desroy et al., 2017). CpdA was synthetized in five steps as depicted below.

##### Synthesis of Int 1

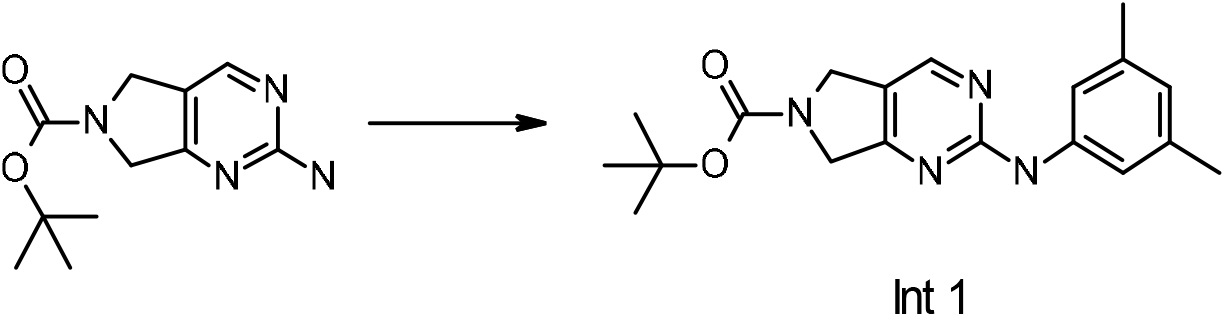

tert-Butyl 2-amino-5h-pyrrolo[3,4-d]pyrimidine-6(7h)-carboxylate (11 g, 46.6 mmol, 1.0 eq.), 1-bromo-3,5-dimethyl-benzene (9 g, 48.9 mmol, 1.05 eq.), Xantphos (5.49 g, 9.31 mmol, 0.2 eq.), NaOtBu (13.8 g, 139.7 mmol, 3.0 eq.), and Pd_2_(dba)_3_ (4.35 g, 4.7 mmol, 0.1 eq.) were placed in 1,4-dioxane (150 mL) under Ar atmosphere and the mixture was vigorously stirred at 110 °C for 3 h then at 80 °C overnight. The reaction medium was poured into a well-stirred mixture of NaCl (250 g) in EtOAc/water (2/1, 1.5 L). The layers were separated, and the aqueous phase was extracted with EtOAc. The combined organic layers were filtered over Celite®, dried over Na_2_SO_4_, filtered, and concentrated. The residue was suspended in an EtOAc/cyclohexane mixture (1/1, 200 mL) and the resulting precipitate was filtered and washed with an EtOAc/cyclohexane mixture (1/1). The solid was dried to afford Int 1.

##### Synthesis of Int 2

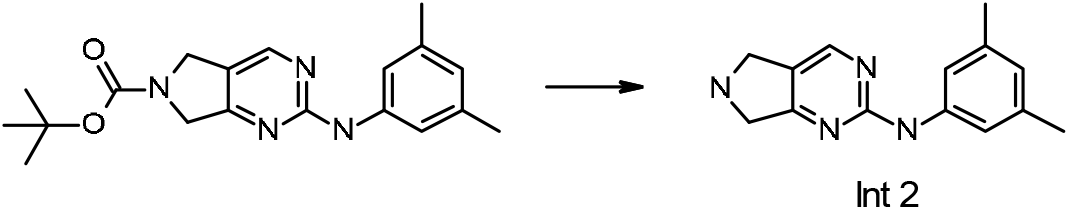

**Int 1** (39.8 g, 1151.7 mmol, 1.0 eq.) was suspended in a solution of 4N HCl in 1,4-dioxane (400 mL). The mixture was stirred at RT for 3 h, then concentrated to dryness. The resulting solid was dissolved in water, and DCM and 2N NaOH were added. The layers were separated and the aqueous layer was back-extracted twice with DCM. The organic layers were evaporated to dryness to obtain **Int 2**.

##### Synthesis of Int 3

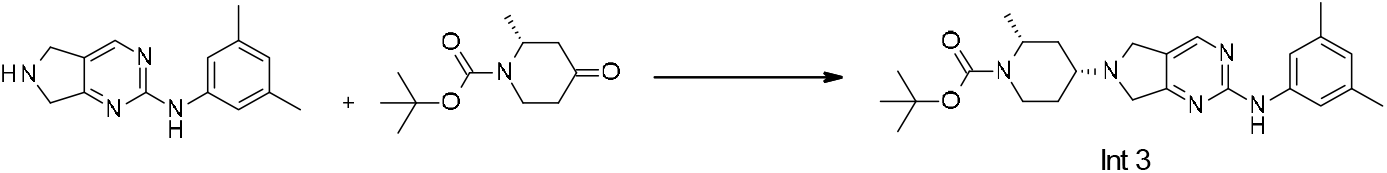

**Int 2** (27 g, 112.4 mmol, 1.0 eq.) and tert-butyl (2R)-2-methyl-4-oxo-piperidine-1-carboxylate (CAS# 790667-43-5; 25.2 g, 118.2 mmol, 1.05 eq.) were stirred in DCM (350 mL) and AcOH (0.7 mL) for 30 min under inert atmosphere. NaBH(OAc)_3_ (35.6 g, 168.0 mmol, 1.5 eq.) was added and the mixture was stirred at RT for 1 h. The mixture was poured in a NaOAc solution before the aqueous phase was extracted with DCM, dried over Na_2_SO_4_, and concentrated to dryness. The crude product was purified by flash column chromatography on silica gel (eluted with heptane/EtOAc) to afford **Int 3**.

##### Synthesis of Int 4

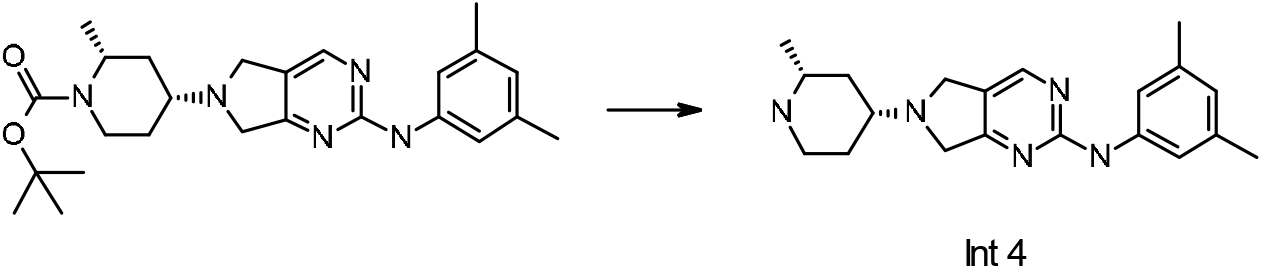

**Int 3** (15.4 g, 35.2 mmol, 1.0 eq.) was suspended in HCl (4N in 1,4-dioxane, 250 mL) and the mixture was stirred at RT for 2 h. The medium was concentrated to dryness and the residue was taken up in DCM. Water was added and the pH of the aqueous phase was adjusted to 10 using 40% NaOH aq. solution. The layers were separated, and the aqueous layer was extracted with DCM. The combined organic phases were concentrated to afford **Int 4**.

##### Synthesis of CpdA

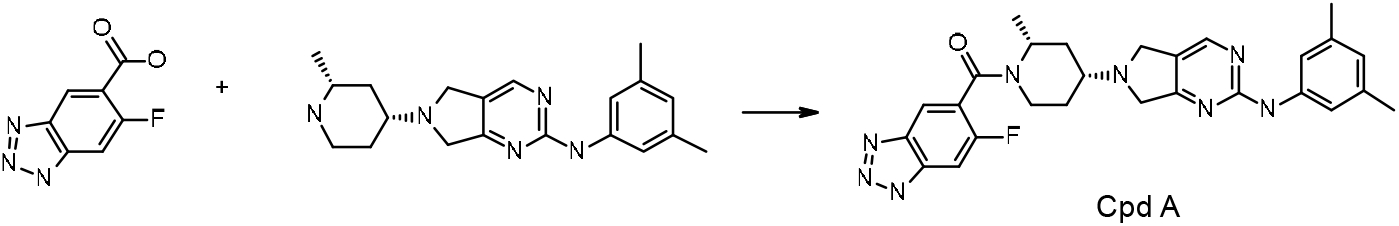

6-fluoro-1H-benzotriazole-5-carboxylic acid (CAS# 1427081-62-6; 5.4 g, 29.8 mmol, 1.0 eq.), HATU (11.4 g, 29.8 mmol, 1.0 eq.), HOAt (2.0 g, 14.9 mmol, 0.5 eq.), and tetramethyl piperidine (10.5 g, 74.5 mmol, 2.5 eq.) were dissolved in DMF (100 mL). The reaction mixture was stirred at RT for 5 min. **Int 4** (10.1 g, 30.0 mmol, 1.03 eq.) was then added and the mixture was stirred at RT overnight. The reaction mixture was poured on a sat. NH_4_Cl aq. solution (400 mL) and stirred for 40 min in an ice bath. The precipitate was filtered off, rinsed with water, dried under suction for 1 h and under vacuum at 40 °C to yield crude material. The crude product was purified by chromatography on silica gel, eluting with DCM:MeOH:NH_4_OH (90:9:1.5). The fractions of interest were pooled and concentrated under reduced pressure. The solid material obtained was triturated in acetone (20 vol.) at 0 °C for 40 min, filtered, and dried under vacuum for 4 h to afford **CpdA**.

### Quantification and statistical analysis

Statistical analysis of experimental results was performed according to the quality standard for each technique, and significance was evaluated using GraphPad 8 software.

### Key resources table

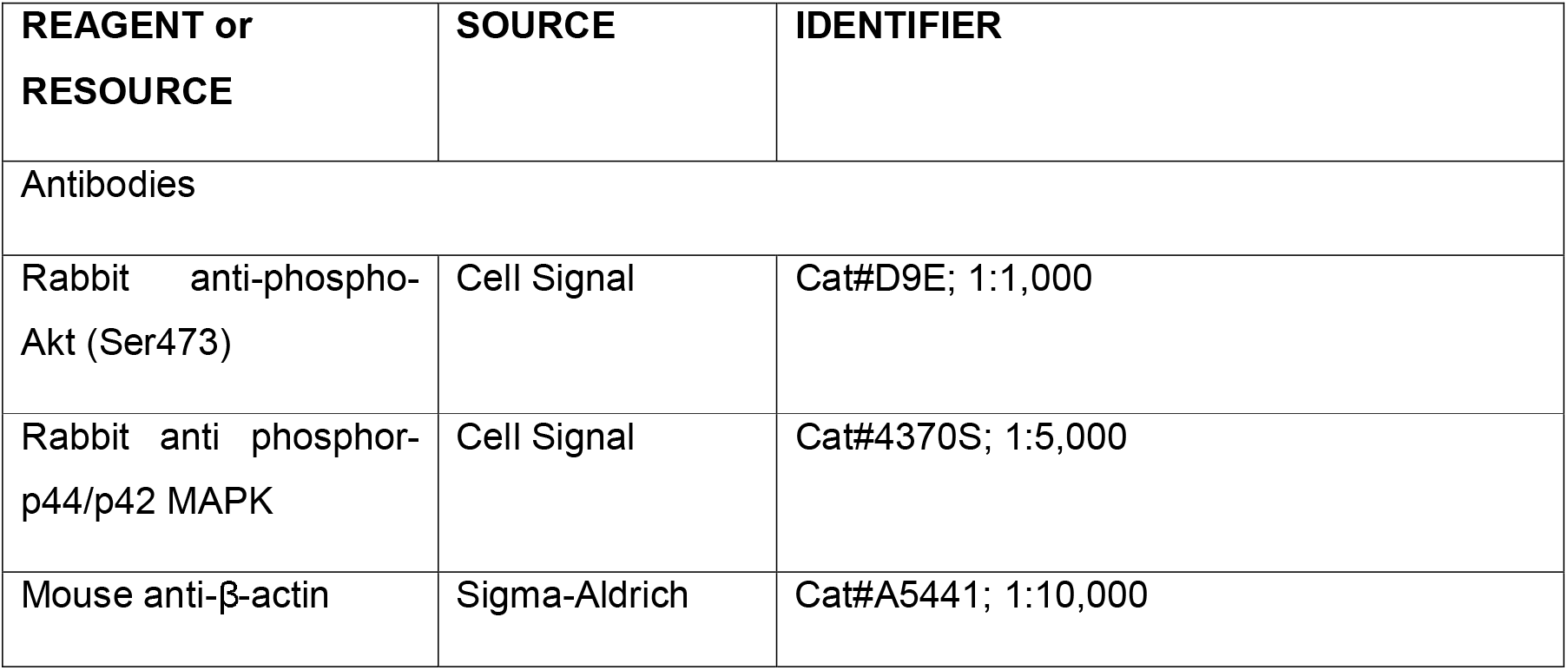
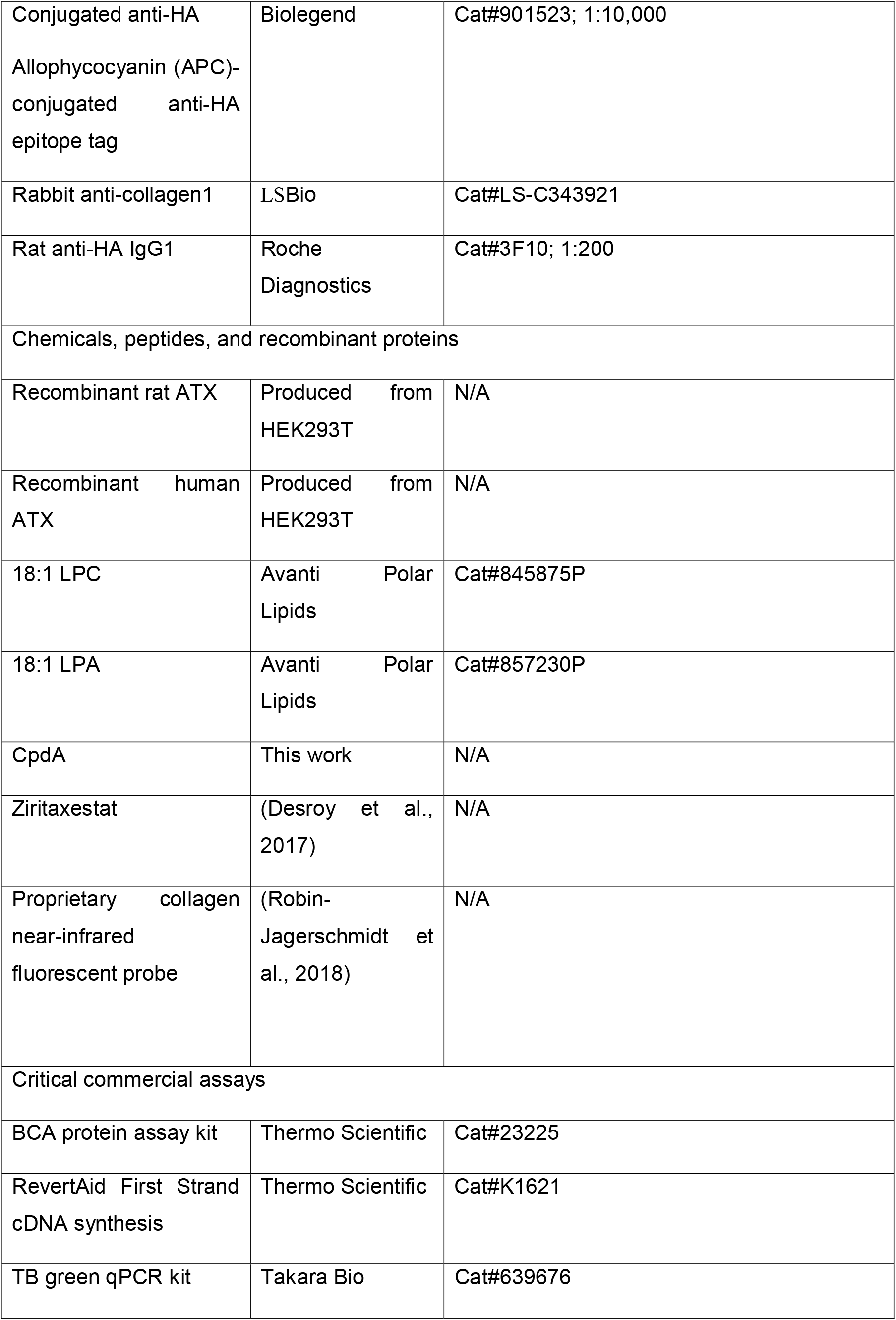
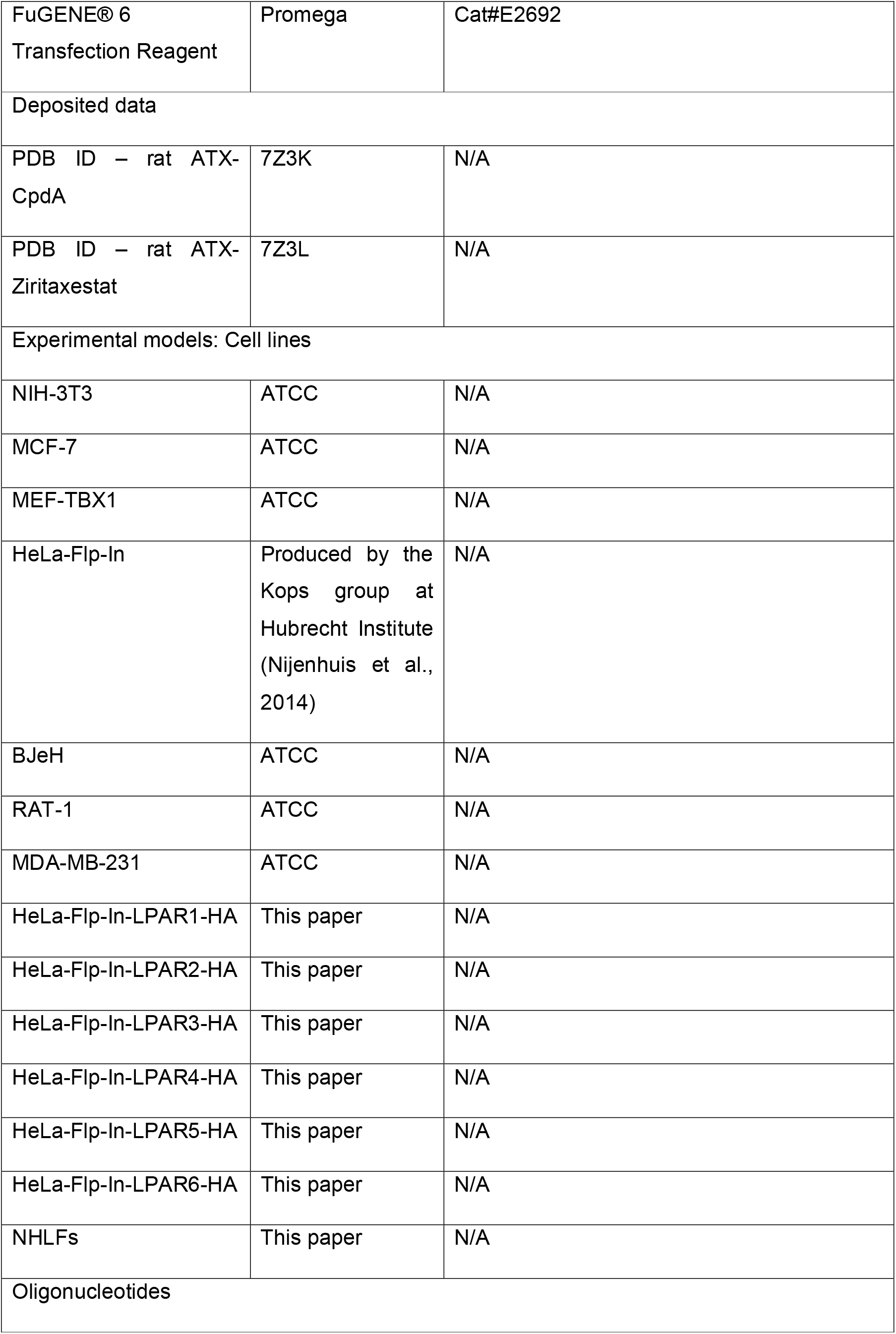
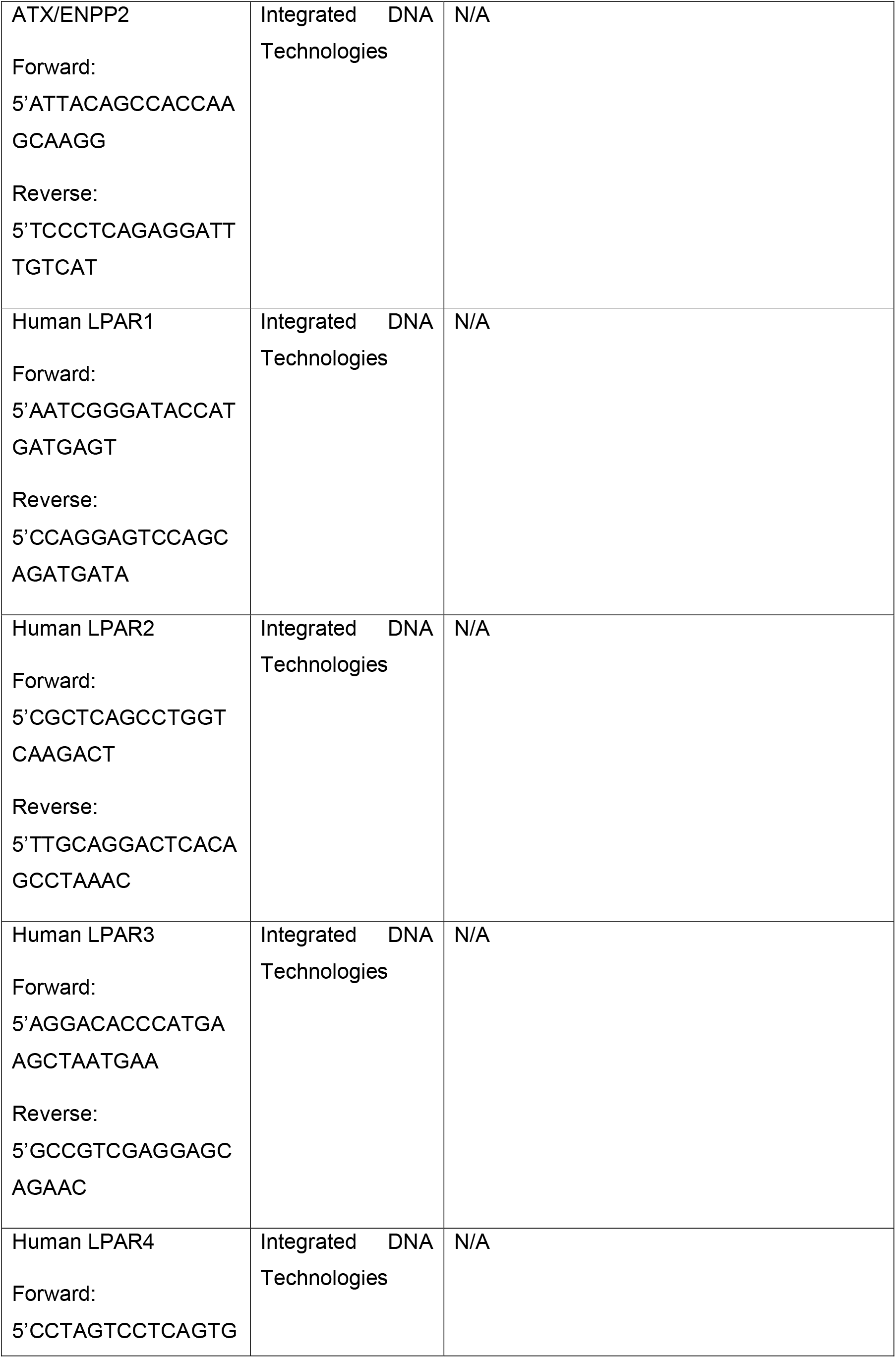
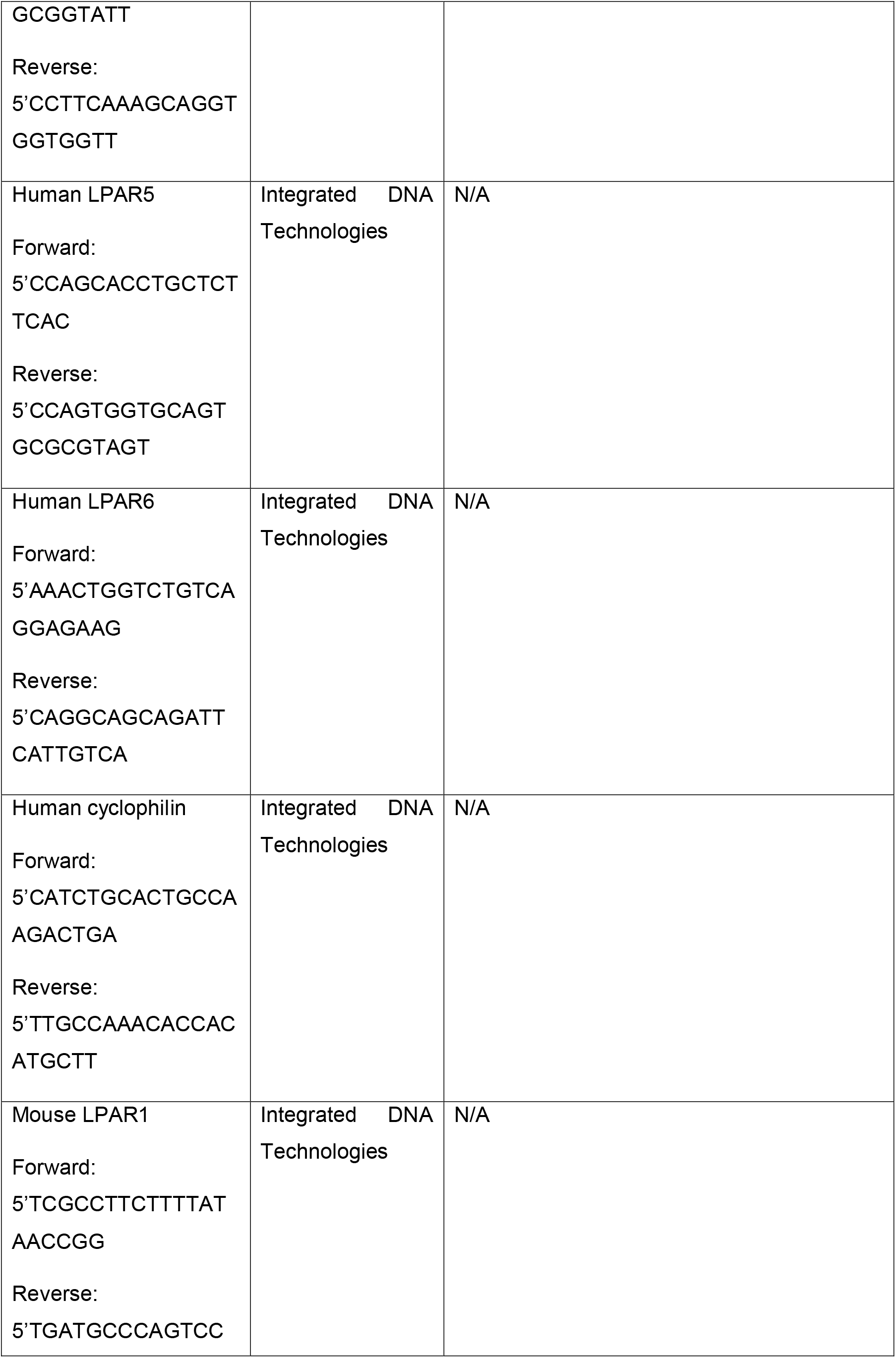
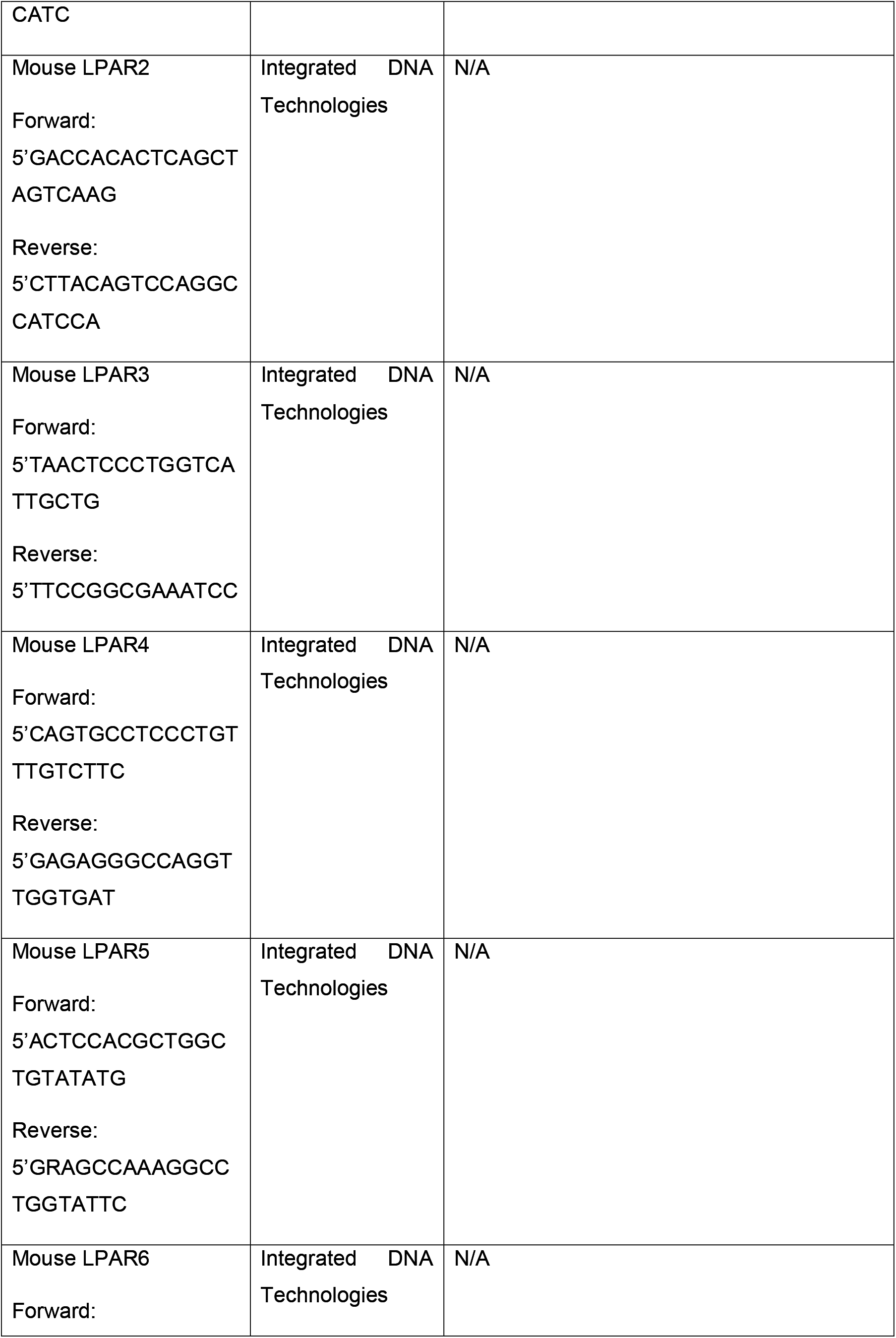
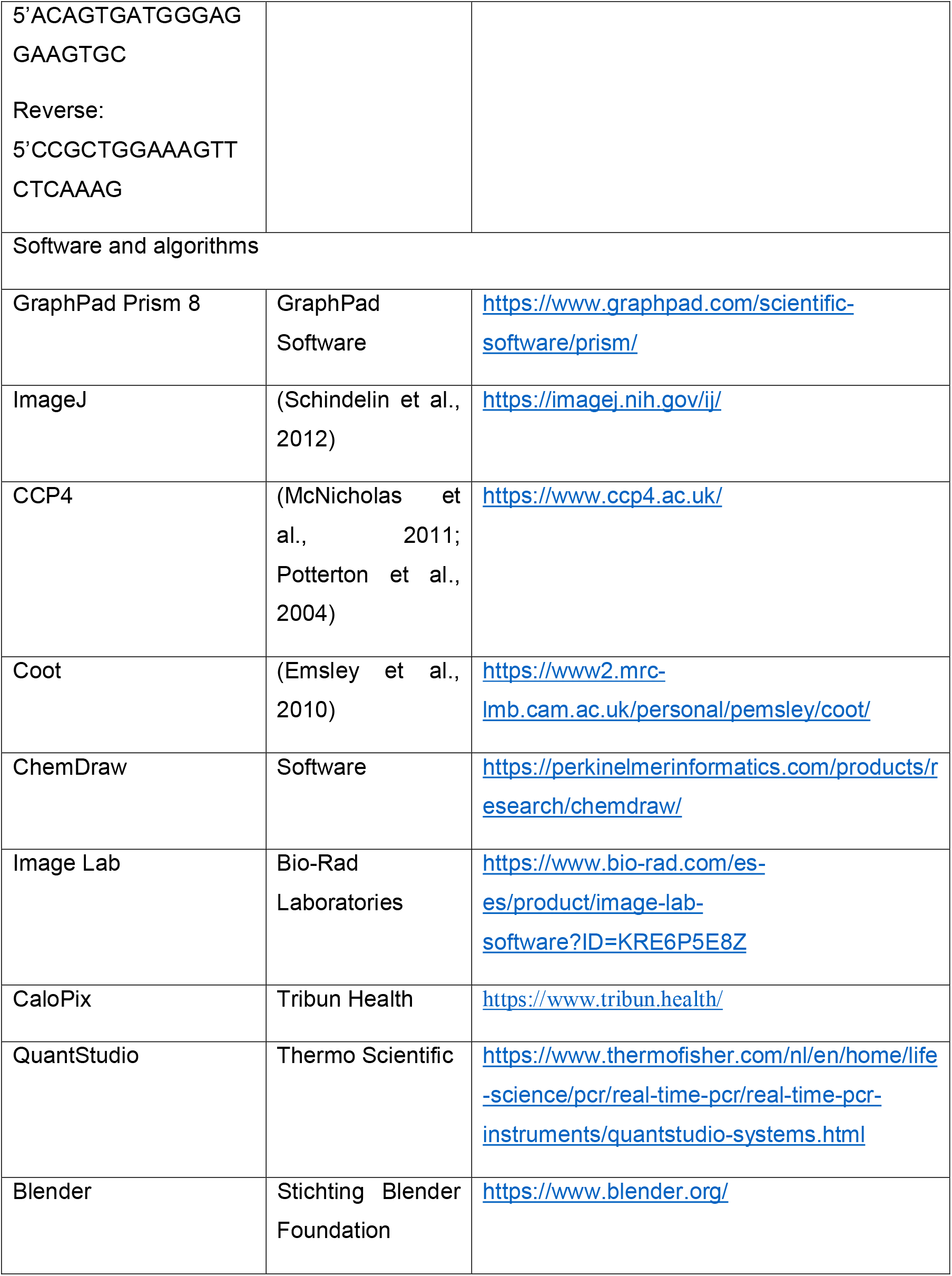

