## Supplementary material for "Autotaxin facilitates selective LPA receptor signaling": Detailed Experimental Procedures

### Supplemental Experimental Procedures

#### Resource availability

**Lead contact**

**Materials availability**

This study did not generate new unique reagents.

**Data and code availability**

The co-crystal structures of ATX bound to CpdA or ziritaxestat were deposited at the Protein Data Bank under the codes 7Z3K and 7Z3L, respectively.

- Crystallographic data have been deposited at the Protein Data Bank and are publicly available as of the date of publication. Accession numbers are listed in the key resources table.
- This paper does not report original code.
- Any additional information required to reanalyze the data reported in this paper is available from the lead contact upon request.

#### Organic chemistry methods

***Structure of GLPG1690 and CpdA***

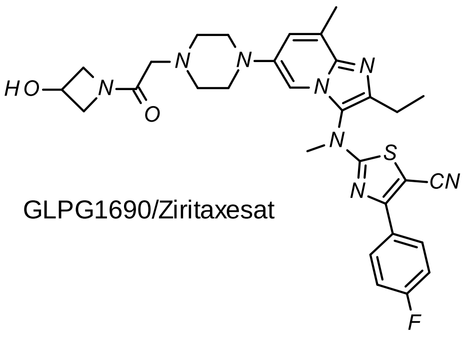


GLPG1690/Ziritaxesat

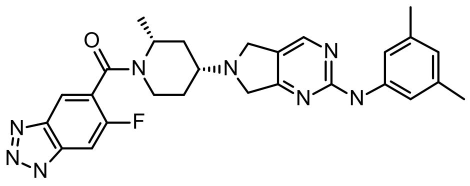


Cpd A

GLPG1690/ziritaxestat synthesis was published in (Desroy et al., 2017). CpdA was synthetized in five steps as depicted below.

***Synthesis of Int 1***

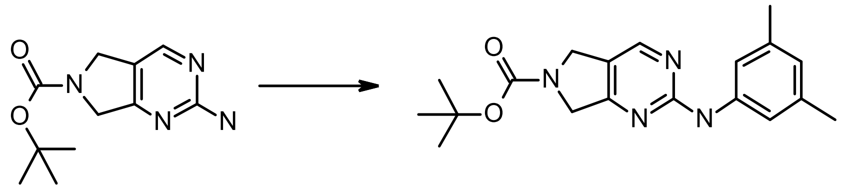

Int 1

tert-Butyl 2-amino-5h-pyrrolo[3,4-d]pyrimidine-6(7h)-carboxylate (11 g, 46.6 mmol, 1.0 eq.), 1-bromo-3,5-dimethyl-benzene (9 g, 48.9 mmol, 1.05 eq.), Xantphos (5.49 g, 9.31 mmol, 0.2 eq.), NaOtBu (13.8 g, 139.7 mmol, 3.0 eq.), and $Pd_2(dba)_3$ (4.35 g, 4.7 mmol, 0.1 eq.) were placed in 1,4-dioxane (150 mL) under Ar atmosphere and the mixture was vigorously stirred at 110 °C for 3 h then at 80 °C overnight. The reaction medium was poured into a well-stirred mixture of NaCl (250 g) in EtOAc/water (2/1, 1.5 L). The layers were separated, and the aqueous phase was extracted with EtOAc. The combined organic layers were filtered over Celite®, dried over $Na_2SO_4$, filtered, and concentrated. The residue was suspended in an EtOAc/cyclohexane mixture (1/1, 200 mL) and the resulting precipitate was filtered and washed with an EtOAc/cyclohexane mixture (1/1). The solid was dried to afford Int 1.

***Synthesis of Int 2***

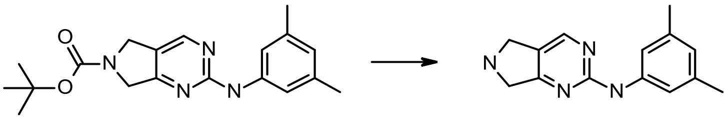

Int 2

**Int 1** (39.8 g, 1151.7 mmol, 1.0 eq.) was suspended in a solution of 4N HCl in 1,4-dioxane (400 mL). The mixture was stirred at RT for 3 h, then concentrated to dryness. The resulting solid was dissolved in water, and DCM and 2N NaOH were added. The layers were separated and the aqueous layer was back-extracted twice with DCM. The organic layers were evaporated to dryness to obtain **Int 2**.

***Synthesis of Int 3***

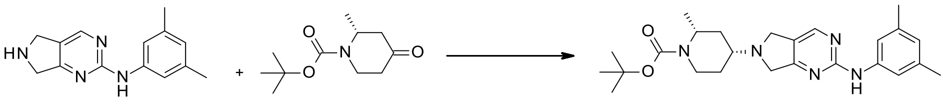

Int 3

**Int 2** (27 g, 112.4 mmol, 1.0 eq.) and tert-butyl (2R)-2-methyl-4-oxo-piperidine-1-carboxylate (CAS# 790667-43-5; 25.2 g, 118.2 mmol, 1.05 eq.) were stirred in DCM (350 mL) and AcOH (0.7 mL) for 30 min under inert atmosphere. $NaBH(OAc)_3$ (35.6 g, 168.0 mmol, 1.5 eq.) was added and the mixture was stirred at RT for 1 h. The mixture was poured in a NaOAc solution before the aqueous phase was extracted with DCM, dried over $Na_2SO_4$, and concentrated to dryness. The crude product was purified by flash column chromatography on silica gel (eluted with heptane/EtOAc) to afford **Int 3**.

***Synthesis of Int 4***

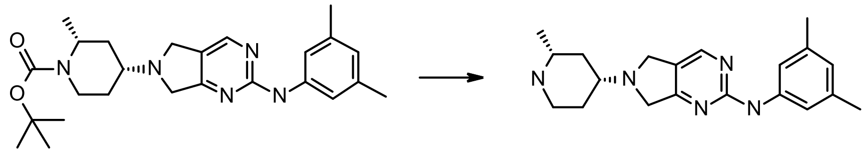

Int 4

**Int 3** (15.4 g, 35.2 mmol, 1.0 eq.) was suspended in HCl (4N in 1,4-dioxane, 250 mL) and the mixture was stirred at RT for 2 h. The medium was concentrated to dryness and the residue was taken up in DCM. Water was added and the pH of the aqueous phase was adjusted to 10 using 40% NaOH aq. solution. The layers were separated, and the aqueous layer was extracted with DCM. The combined organic phases were concentrated to afford **Int 4**.

***Synthesis of CpdA***

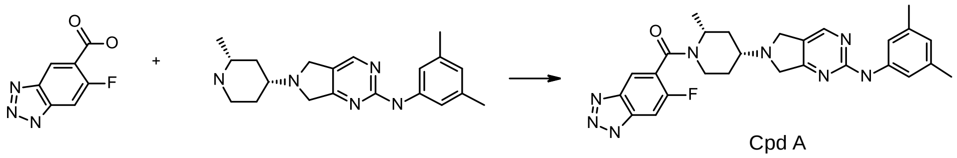

Cpd A

6-fluoro-1H-benzotriazole-5-carboxylic acid (CAS# 1427081-62-6; 5.4 g, 29.8 mmol, 1.0 eq.), HATU (11.4 g, 29.8 mmol, 1.0 eq.), HOAt (2.0 g, 14.9 mmol, 0.5 eq.), and tetramethyl piperidine (10.5 g, 74.5 mmol, 2.5 eq.) were dissolved in DMF (100 mL). The reaction mixture was stirred at RT for 5 min. **Int 4** (10.1 g, 30.0 mmol, 1.03 eq.) was then added and the mixture was stirred at RT overnight. The reaction mixture was poured on a sat. $NH_4Cl$ aq. solution (400 mL) and stirred for 40 min in an ice bath. The precipitate was filtered off, rinsed with water, dried under suction for 1 h and under vacuum at 40 °C to yield crude material. The crude product was purified by chromatography on silica gel, eluting with DCM:MeOH:$NH_4OH$ (90:9:1.5). The fractions of interest were pooled and concentrated under reduced pressure. The solid material obtained was triturated in acetone (20 vol.) at 0 °C for 40 min, filtered, and dried under vacuum for 4 h to afford **CpdA**.

#### Quantification and statistical analysis

Statistical analysis of experimental results was performed according to the quality standard for each technique, and significance was evaluated using GraphPad 8 software.

### Key resources table

| REAGENT or RESOURCE | SOURCE | IDENTIFIER |
|---|---|---|
| Antibodies | | |
| Rabbit anti-phospho-Akt (Ser473) | Cell Signal | Cat#D9E; 1:1,000 |
| Rabbit anti phosphor-p44/p42 MAPK | Cell Signal | Cat#4370S; 1:5,000 |
| Mouse anti-β-actin | Sigma-Aldrich | Cat#A5441; 1:10,000 |
| Conjugated anti-HA<br><br>Allophycocyanin (APC)-conjugated anti-HA epitope tag | Biolegend | Cat#901523; 1:10,000 |
| Rabbit anti-collagen1 | LSBio | Cat#LS-C343921 |
| Rat anti-HA IgG1 | Roche Diagnostics | Cat#3F10; 1:200 |
| Chemicals, peptides, and recombinant proteins | | |
| Recombinant rat ATX | Produced from HEK293T | N/A |
| Recombinant human ATX | Produced from HEK293T | N/A |
| 18:1 LPC | Avanti Polar Lipids | Cat#845875P |
| 18:1 LPA | Avanti Polar Lipids | Cat#857230P |
| CpdA | This work | N/A |
| Ziritaxestat | (Desroy et al., 2017) | N/A |

| | | |
|---|---|---|
| Proprietary collagen near-infrared fluorescent probe | (Robin-Jagerschmidt et al., 2018) | N/A |
| Critical commercial assays | | |
| BCA protein assay kit | Thermo Scientific | Cat#23225 |
| RevertAid First Strand cDNA synthesis | Thermo Scientific | Cat#K1621 |
| TB green qPCR kit | Takara Bio | Cat#639676 |
| FuGENE® 6 Transfection Reagent | Promega | Cat#E2692 |
| Deposited data | | |
| PDB ID – rat ATX-CpdA | 7Z3K | N/A |
| PDB ID – rat ATX-Ziritaxestat | 7Z3L | N/A |
| Experimental models: Cell lines | | |
| NIH-3T3 | ATCC | N/A |
| MCF-7 | ATCC | N/A |
| MEF-TBX1 | ATCC | N/A |
| HeLa-Flp-In | Produced by the Kops group at Hubrecht Institute (Nijenhuis et al., 2014) | N/A |
| BJeH | ATCC | N/A |
| RAT-1 | ATCC | N/A |
| MDA-MB-231 | ATCC | N/A |
| HeLa-Flp-In-LPAR1-HA | This paper | N/A |

| | | |
|---|---|---|
| HeLa-Flp-In-LPAR2-HA | This paper | N/A |
| HeLa-Flp-In-LPAR3-HA | This paper | N/A |
| HeLa-Flp-In-LPAR4-HA | This paper | N/A |
| HeLa-Flp-In-LPAR5-HA | This paper | N/A |
| HeLa-Flp-In-LPAR6-HA | This paper | N/A |
| NHLFs | This paper | N/A |
| Oligonucleotides | | |
| ATX/ENPP2<br>Forward: 5'ATTACAGCCACCAAGCAAGG<br>Reverse: 5'TCCCTCAGAGGATTTGTCAT | Integrated DNA Technologies | N/A |
| Human LPAR1<br>Forward: 5'AATCGGGATACCATGATGAGT<br>Reverse: 5'CCAGGAGTCCAGCAGATGATA | Integrated DNA Technologies | N/A |
| Human LPAR2<br>Forward: 5'CGCTCAGCCTGGTCAAGACT<br>Reverse: 5'TTGCAGGACTCACAGCCTAAAC | Integrated DNA Technologies | N/A |

| | | |
|---|---|---|
| Human LPAR3<br>Forward: 5’AGGACACCCATGAAGCTAATGAA<br>Reverse: 5’GCCGTCGAGGAGCAGAAC | Integrated DNA Technologies | N/A |
| Human LPAR4<br>Forward: 5’CCTAGTCCTCAGTGGCGGTATT<br>Reverse: 5’CCTTCAAAGCAGGTGGTGGTT | Integrated DNA Technologies | N/A |
| Human LPAR5<br>Forward: 5’CCAGCACCTGCTCTTCAC<br>Reverse: 5’CCAGTGGTGCAGTGCGCGTAGT | Integrated DNA Technologies | N/A |
| Human LPAR6<br>Forward: 5’AAACTGGTCTGTCAGGAGAAG<br>Reverse: 5’CAGGCAGCAGATTCATTGTCA | Integrated DNA Technologies | N/A |
| Human cyclophilin | Integrated DNA Technologies | N/A |

| | | |
|---|---|---|
| Forward: 5’CATCTGCACTGCCAAGACTGA<br><br>Reverse: 5’TTGCCAAACACCACATGCTT | | |
| Mouse LPAR1<br><br>Forward: 5’TCGCCTTCTTTTATAACCGG<br><br>Reverse: 5’TGATGCCCAGTCCCATC | Integrated DNA Technologies | N/A |
| Mouse LPAR2<br><br>Forward: 5’GACCACACTCAGCTAGTCAAG<br><br>Reverse: 5’CTTACAGTCCAGGCCATCCA | Integrated DNA Technologies | N/A |
| Mouse LPAR3<br><br>Forward: 5’TAACTCCCTGGTCATTGCTG<br><br>Reverse: 5’TTCCGGCGAAATCC | Integrated DNA Technologies | N/A |
| Mouse LPAR4<br><br>Forward: 5’CAGTGCCTCCCTGTTTGTCTTC | Integrated DNA Technologies | N/A |

| | | |
|---|---|---|
| Reverse: 5'GAGAGGGCCAGGTTGGTGAT | | |
| Mouse LPAR5<br><br>Forward: 5'ACTCCACGCTGGCTGTATATG<br><br>Reverse: 5'GRAGCCAAAGGCCTGGTATTC | Integrated DNA Technologies | N/A |
| Mouse LPAR6<br><br>Forward: 5'ACAGTGATGGGAGGAAGTGC<br><br>Reverse: 5'CCGCTGGAAAGTTCTCAAAG | Integrated DNA Technologies | N/A |
| Software and algorithms | | |
| GraphPad Prism 8 | GraphPad Software | https://www.graphpad.com/scientific-software/prism/ |
| ImageJ | (Schindelin et al., 2012) | https://imagej.nih.gov/ij/ |
| CCP4 | (McNicholas et al., 2011; Potterton et al., 2004) | https://www.ccp4.ac.uk/ |
| Coot | (Emsley et al., 2010) | https://www2.mrc-lmb.cam.ac.uk/personal/pemsley/coot/ |
| ChemDraw | Software | https://perkinelmerinformatics.com/products/research/chemdraw/ |

| | | |
|---|---|---|
| Image Lab | Bio-Rad Laboratories | https://www.bio-rad.com/es-es/product/image-lab-software?ID=KRE6P5E8Z |
| CaloPix | Tribun Health | https://www.tribun.health/ |
| QuantStudio | Thermo Scientific | https://www.thermofisher.com/nl/en/home/life-science/pcr/real-time-pcr/real-time-pcr-instruments/quantstudio-systems.html |
| Blender | Stichting Blender Foundation | https://www.blender.org/ |
