## Supplementary material for "Autotaxin facilitates selective LPA receptor signaling": Fig. S

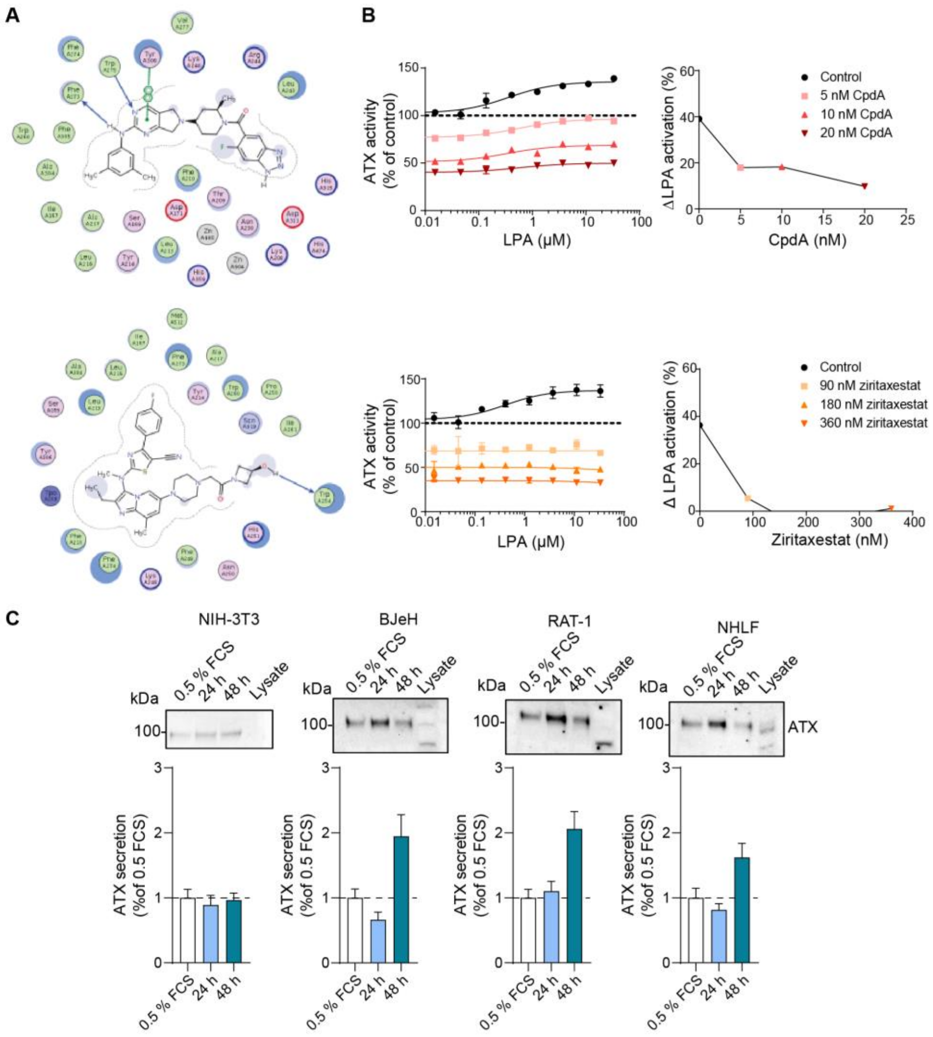


**Figure S1. Different types of ATX inhibitors modulate specific signaling events**.

(**A**) Interaction of CpdA (upper panel) and ziritaxestat (lower panel) with ATX, shown in a two-dimensional interaction network generated by Lidia software on WinCoot.

(**B**) Ability of CpdA (upper panels) and ziritaxestat (lower panels) to block the ATX tunnel, determined by the abolition of LPA allostery while retaining residual, slow-turnover ATX catalysis. The dashed horizontal line represents the time of cell stimulation. Data represent the average value of triplicate measures ± SEM (error bars).

(**C**) Secretion of ATX by the cell lines evaluated in this study. Data represent the average value of triplicate measures ± SEM (error bars).

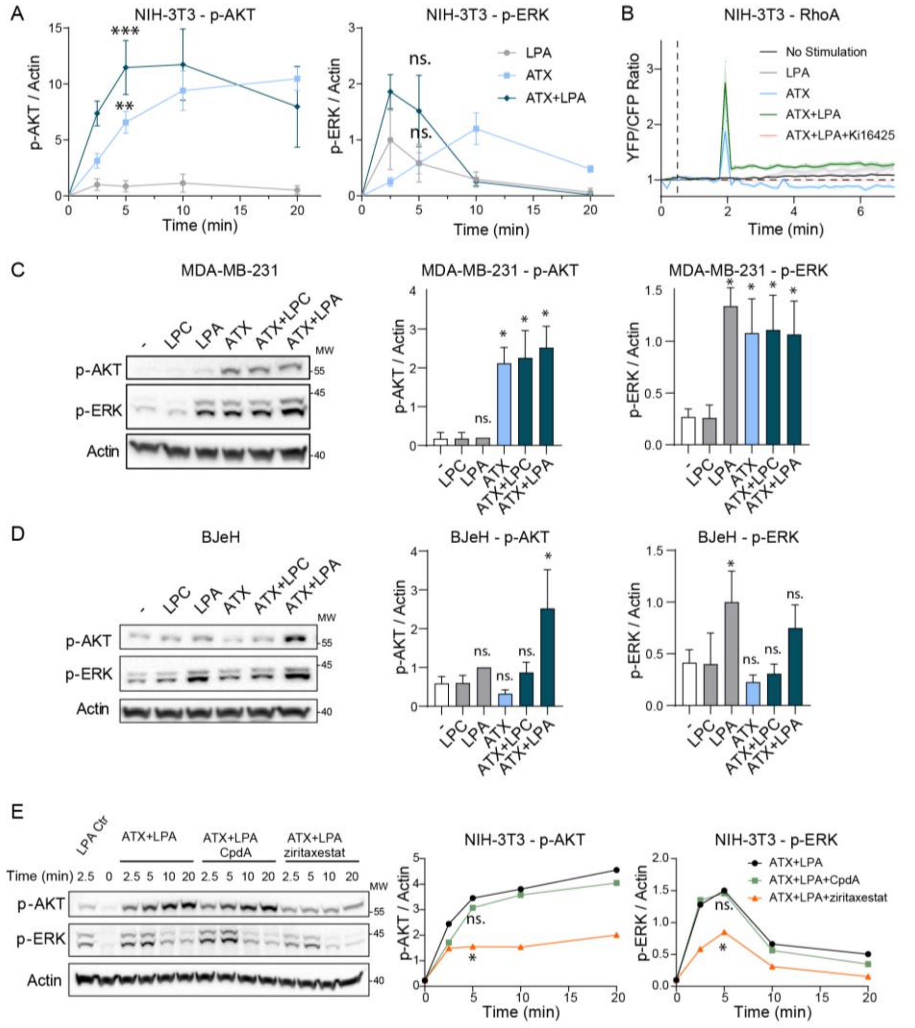


**Figure S2. ATX is crucial for triggering LPA signaling**.

(**A**) Kinetics of NIH-3T3 cell responses through AKT and ERK activation normalized to LPA control after a 2.5-min stimulation. Quantitation of three independent experiments, shown as the mean ± SEM. **p<0.01; ***p<0.001; ns, not significant (one-way ANOVA).

(**B**) Raw data behind the YFP/CFP fluorescence ratio shown in **Fig. 2B**. The dashed vertical line represents the time of cell stimulation. The ratio was calculated upon time-course stimulation of NIH-3T3 cells with LPA, ATX, or ATX-bound LPA for 10 min in the presence or absence of 5 μM CpdA or ziritaxestat.

(**C,D**) Activation of AKT and ERK in response to LPC, LPA, ATX, and the combination thereof in (**C**) MDA-MB-231 and (**D**) BJeH cells. ATX was preincubated with LPC or LPA for at least 30 min. Quantitation of three independent experiments, shown as the mean ± SEM. *p<0.05; ns, not significant (one-way ANOVA).

(**E**) Blockade of the activation of AKT and ERK over time by 5 μM CpdA and ziritaxestat in NIH-3T3 cells. Quantitation of three independent experiments, shown as the mean ± SEM. *p<0.05; ns, not significant (one-way ANOVA).

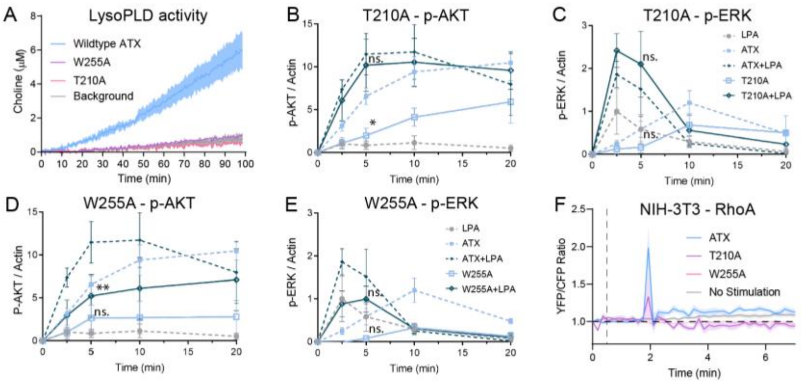


**Figure S3. LPA delivery is catalysis independent and requires an intact ATX tunnel**.

(**A**) Choline release assays of wildtype ATX, catalytically inactive ATX(T210A), and tunnel mutant ATX(W255A). Data represent the average of three different experiments ± SD.

(**B–E**) Activation of AKT and ERK in NIH-3T3 cells over time upon stimulation with LPA and (**B,C**) ATX(T210A) or (**D,E**) ATX(W255A), or LPA bound to each mutant. Quantitation of three independent experiments, shown as the mean ± SEM. *p<0.05; **p<0.01; ns, not significant (one-way ANOVA).

(**F**) Complete time course measure of RhoA activation as YFP/CFP fluorescence ratio from quantitation in **Fig. 3**. The dashed vertical line represents the time of cell stimulation.

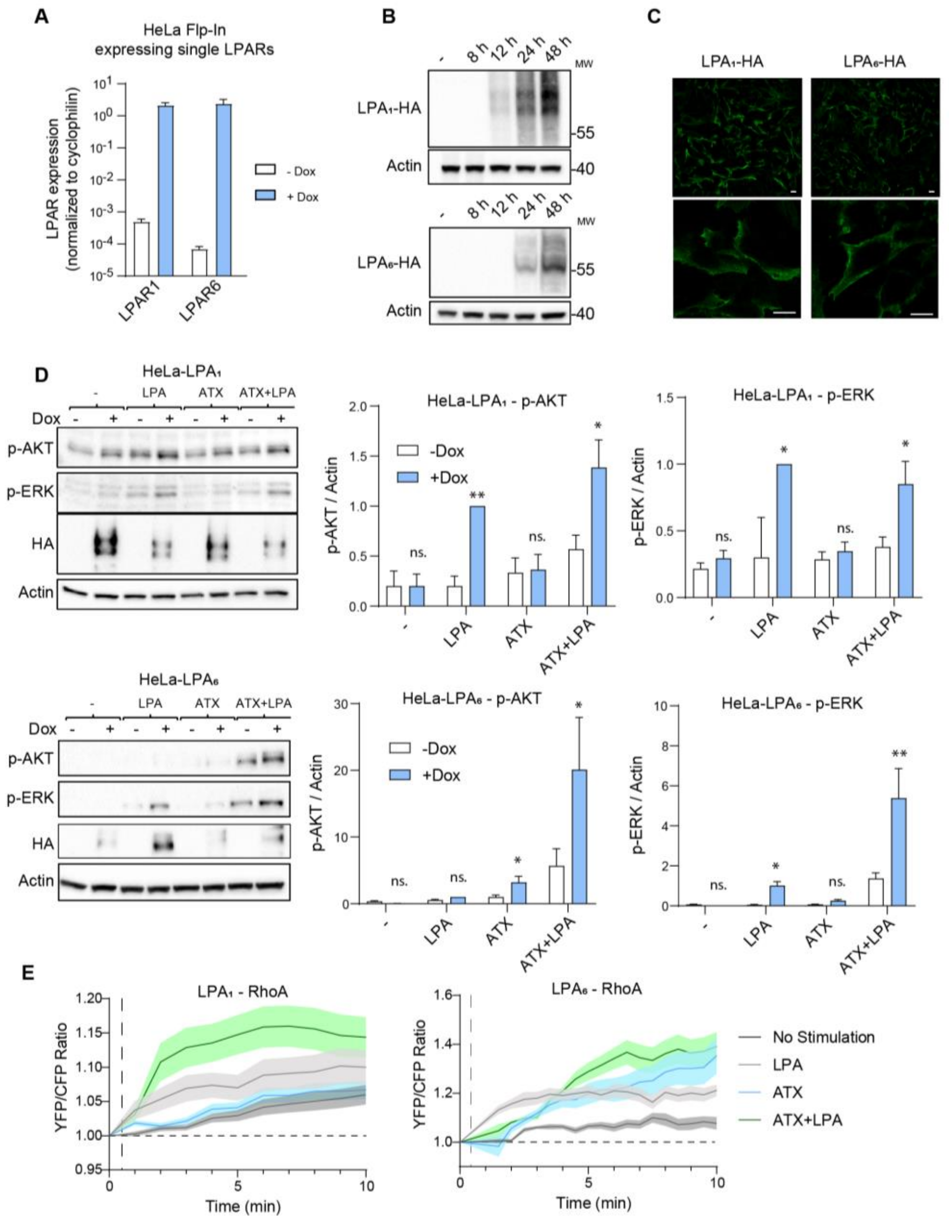


**Figure S4. ATX-mediated signaling favors P2Y LPA$_6$ receptor**.

(**A**) qPCR expression of induced HA-tagged LPARs compared with that of uninduced HeLa-Flp-In cells. Ct values were normalized to cyclophilin and presented in logarithmic scale.

(**B**) Representative Western blots and quantitation of uninduced and induced LPA$_1$-HA- and LPA$_6$-HA-expressing HeLa-Flp-In cells. Quantitation of three independent experiments, representing the average value of triplicate biological measures ± SEM (error bars).

(**C**) Representative confocal microscopy images of LPAR induction and subcellular localization at the plasma membrane. Scale bar represents 10 μm.

(**D**) Stimulation of HeLa-Flp-In cells that were starved overnight with 0.5% serum-containing medium, where receptor expression was also induced. Left panels, representative Western blots of AKT and ERK activation; center and right panels, quantitation of p-AKT and p-ERK from three independent experiments, shown as the mean ± SEM. *p<0.05, **p<0.01; ns, not significant (one-way ANOVA).

(**E**) Complete time-course measure of RhoA activation as YFP/CFP fluorescence ratio from quantitation in **Fig. 4**. The dashed vertical line represents the time of cell stimulation.

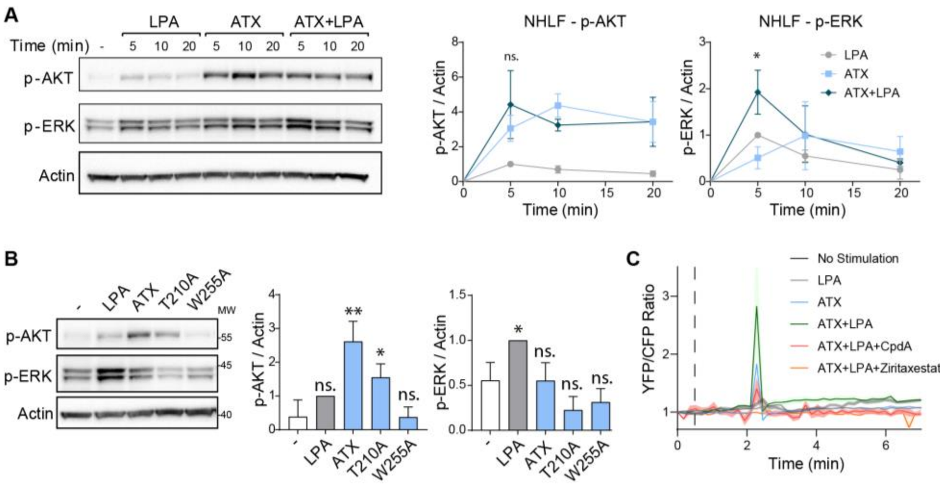


**Figure S5. ATX-mediated LPA delivery and signaling in primary lung fibroblasts**.

(**A**) NHLFs were serum starved for 16 h and stimulated at different times with LPA (1 μM), ATX (20 nM), or a mixture of both. Upper panel, representative Western blot; lower panels, fold-change of AKT and ERK activation kinetics. Quantitation of three independent experiments, representing the average value of triplicate biological measures ± SEM (error bars). *p<0.05; ns, not significant (unpaired t-test).

(**B**) NHLFs were stimulated with LPA, ATX, catalytically inactive ATX(T210), and tunnel mutant ATX(W255A), and analyzed by Western blotting. Left panel, representative Western blots of AKT and ERK activation (total AKT, total ERK, and actin are shown as loading controls); right panels, quantitation of p-AKT and p-ERK from three independent experiments, representing the average value of triplicate biological measures ± SEM (error bars). *p<0.05, **p<0.01; ns, not significant (unpaired t-test).

(**C**) Complete time-course measure of RhoA activation as YFP/CFP fluorescence ratio from quantitation in **Fig. 5**. The dashed vertical line represents the time of cell stimulation.

Steady state Pharmacokinetic parameters

| Plasma parameters | | Ziritaxestat | CpdA |
|---|---|---|---|
| Female albino C57BL/6J mice | | 30 mg/kg p.o. b.i.d. MC 0.5 % | 10 mg/kg p.o. b.i.d. PEG400/MC 0.5 % (20/80) |
| **Cmax** | (ng/mL) | 16,600 | 3,430 |
| **Tmax** | (h) | 1.5 | 1 |
| **$C_{last}$** | (ng/mL) | 488 | 8.37 |
| **AUC (0-6h)** | (ng h/mL) | 42,400 | 11,000 |
| **AUC (0-18h)** | (ng h/mL) | 53,500 | 13,200 |
| **AUC (0-24h)** | (ng h/mL) | 95,900 | 24,200 |
| **Rat LPA assay $IC_{50}$** | (nM) | 541 | 13.6 |
| | (ng/mL) | 318 | 7 |

AUC(0- 24 h) = AUX(0-6 h)+AUC(0-18 h)

**Table S1. Ziritaxestat, but not CpdA, reverses pulmonary fibrosis *in vivo*.**

Steady-state pharmacokinetic parameters of CpdA and ziritaxestat.
